# REM Sleep Disengagement of β Oscillations Permits Rapid Dream Movements in Parkinsonism

**DOI:** 10.64898/2026.08.10.743926

**Authors:** Xiaowei Liu, Jing Guang, Zvi Israel, Denise Wajnsztajn, Aeyal Raz, Hagai Bergman

**Author notes:** These authors contributed equally (Xiaowei Liu, Jing Guang). Corresponding authors: Xiaowei Liu, Jing Guang, and Hagai Bergman.;.

## Abstract

REM sleep behavior disorder is a hallmark of prodromal α-synucleinopathies, yet why patients with Parkinson’s disease can generate rapid, coordinated movements during REM sleep despite daytime bradykinesia remains unknown. Here we combined recordings of eye movements, cortical electroencephalography, and basal ganglia-thalamic neuronal activity across vigilance states in non-human primates before and after MPTP-induced parkinsonism. Parkinsonism enhanced β oscillations and impaired movement-related neural dynamics during wakefulness and NREM sleep, whereas both pathological β activity and motor impairments were attenuated during REM sleep. β suppression preceded the onset of REM sleep, indicating that network reconfiguration begins before REM becomes behaviorally apparent. These findings demonstrate that Parkinsonian network dysfunction is dynamically gated by brain state rather than continuously imposed by dopamine depletion. REM sleep is a natural physiological condition in which β oscillations are disengaged, uncovering a latent capacity, normally masked by REM atonia, for rapid movement even in the dopamine-depleted motor network.

## Introduction

Bradykinesia and akinesia are cardinal motor features of Parkinson’s disease (PD) and are closely associated with excessive beta-band (13–30 Hz) synchronized oscillations throughout the cortico-basal ganglia-thalamic network. Beta oscillations correlate with motor impairment, are suppressed by dopaminergic therapy and deep brain stimulation (DBS), and are widely regarded as a physiological hallmark of the Parkinsonian state(*1*). This framework implicitly assumes that pathological beta oscillations, and the resulting motor impairment, are continuously expressed in the dopamine-depleted brain. However, whether parkinsonian network dysfunction is invariant across all naturally occurring brain states remains unknown.

Rapid eye movement (REM, paradoxical) sleep presents a striking challenge to this view. REM sleep is characterized by cortical activation, rapid eye movements, and generalized skeletal muscle atonia generated by brainstem circuits(*2–4*). In patients with REM sleep behavior disorder (RBD), degeneration of these brainstem circuits abolishes REM atonia, allowing dream-enactment behaviors to emerge(*5*, *6*). Because RBD frequently precedes the onset of PD and other synucleinopathies, it has become one of the strongest prodromal markers of neurodegeneration(*7*). Remarkably, despite severe bradykinesia during wakefulness, PD patients with RBD (PD-RBD) can generate rapid, vigorous, and well-coordinated movements and speech during REM sleep(*8*). This paradox suggests that the neural mechanisms responsible for parkinsonian motor impairment are not uniformly expressed across vigilance states.

Although pathological beta activity has been extensively studied during wakefulness, considerably less is known about its dynamics during sleep. Recent recordings have shown that beta oscillations persist during non-rapid eye movement (NREM) sleep and interact with sleep rhythms(*9*, *10*). Yet it remains unknown whether pathological beta spiking activity is maintained during REM sleep, how it evolves during transitions between NREM and REM states, or whether movement-related beta modulation is preserved during REM. Addressing these questions requires simultaneous monitoring of behavior and neuronal activity of multiple brain structures across natural sleep-wake cycles. Here, we combined longitudinal polysomnographic recordings with simultaneous bilateral cortical and subcortical multisite electrophysiological recordings from individual basal ganglia or thalamic nuclei in non-human primates (NHPs) to investigate the relationships between neuronal activity and movement across wakefulness, NREM sleep, and REM sleep before and after MPTP-induced Parkinsonism.

## Results

To characterize neuronal activity across sleep-wake states before and after MPTP, we performed simultaneous polysomnographic and electrophysiological recordings in two African green monkeys during overnight natural sleep (Figs. 1, 2, and S1). Bilateral frontal electroencephalogram (EEG), trapezius electromyography (EMG), and eye movements were recorded simultaneously with bilateral local field potential (LFP) and multi-unit spiking (SPK) activity from either the basal ganglia (GPe and SNr) or thalamic nuclei (VA and CM) (Figs. 1-3 and Table 1). Results from the basal ganglia (BG) and thalamic nuclei are presented in aggregated form in the main text, whereas nucleus-specific analyses are provided in the Supplementary Information.

**Figure 1.**
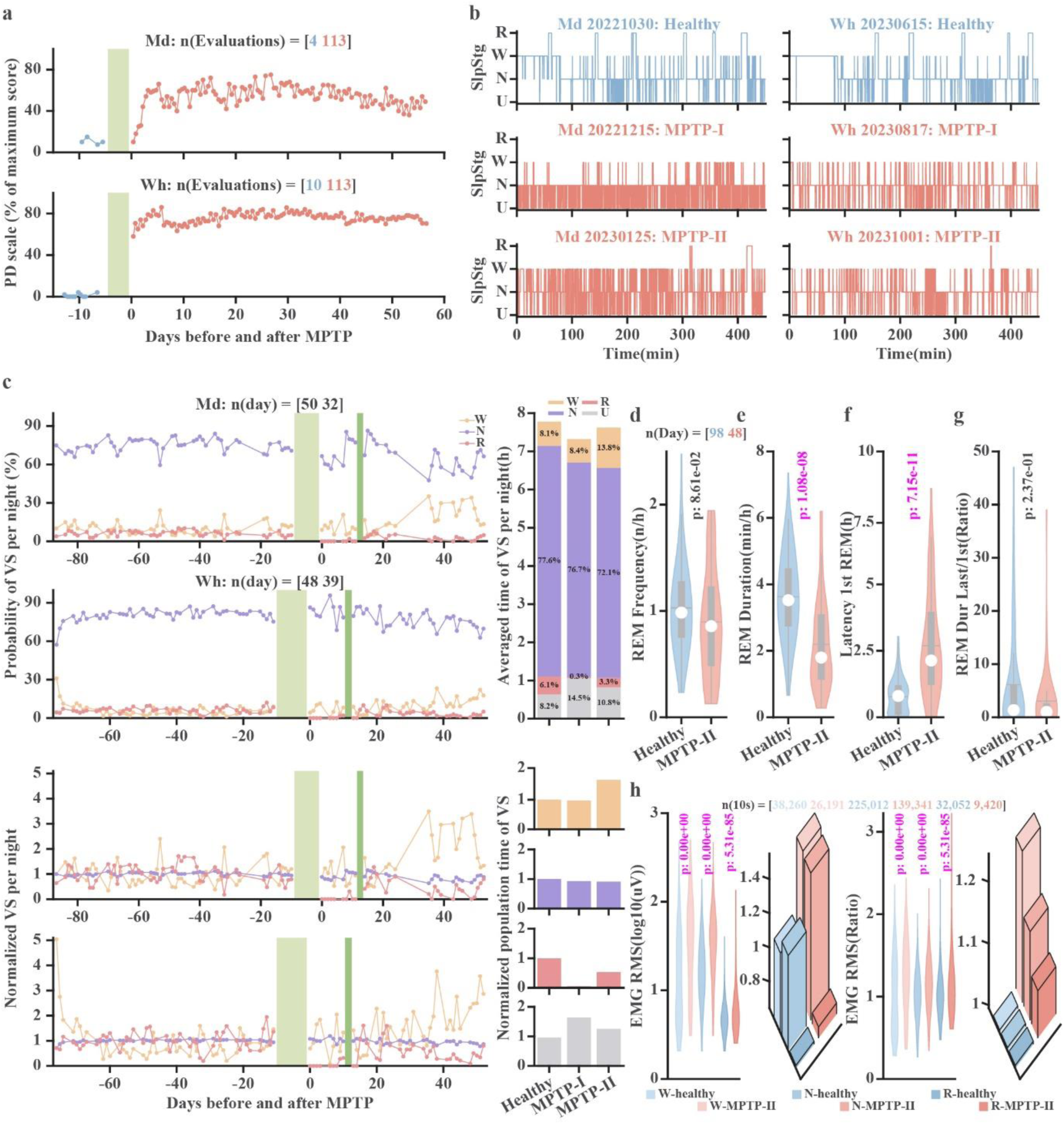
Reduction of REM sleep accompanies MPTP-induced Parkinsonian symptoms across wake–sleep states. **a.** The percentage (out of maximum score) of the PD motor score of the two non-human primates (Md and Wh). The score for each day is the average percentage of all tests on that day. The score evaluated before and after MPTP is separated by a vertical light green bar. Zero indicates the first day after MPTP (2022-12-11 and 2023-08-14 for Md and Wh, respectively). **b.** Examples of hypnograms in healthy (blue), MPTP-I, and MPTP-II (red) states. For comparability, those states are shown in the same time window. **c.** The percentage of wakefulness (yellow), NREM (purple), and REM (pink) per night is shown in the upper-left two rows for Md and Wh, respectively. The population probability of different vigilance stages over the total recording time for the entire night and for both monkeys is shown in the upper-right bar plots. The relative proportions of each state are normalized by their mean probability before MPTP (two bottom-left rows). The normalized population results are shown in the bottom-right bar plots. The vertical light green bar separates the data before and after MPTP. The post-MPTP data is divided (vertical green bar) into MPTP-I and MPTP-II stages (without and with REM episodes, respectively). The meaning of zero is the same as that in **a**. **d-f.** The distribution of REM features before and after MPTP (blue and red, respectively). The violin plots show the mean (grey horizontal lines), median (white circles), 25th and 75th percentiles (the bottom and top edges of the grey boxes, respectively), and the spread of data (grey whiskers). **d.** The number of REM episodes per hour. **e.** REM duration per hour. **f**. Latency to the first REM sleep. **g.** The ratio of the last REM duration to the first REM duration per night. **h.** The distribution of EMG RMS across vigilance states before and after MPTP (first subplot) and a 3-D bar plot of their average values (second subplot). The EMG RMS after MPTP, normalized by the RMS before MPTP, is shown in the third and fourth subplots. The number of units is shown in the title of each subplot. The number of days in **e**-**g** is the same as that in **d**. The statistical differences between variables before and after MPTP in the **d**-**h** subplots are calculated using the Wilcoxon rank-sum test (p<0.05). The p-values indicating significant differences are marked in magenta; otherwise, they are black. Abbreviations. W, wakefulness; N, NREM, non-rapid eye movement; R, REM, rapid eye movement; U, unclassified or unavailable; VS, vigilance states. In the following figures, MPTP denotes the second phase of MPTP (MPTP-II).

**Figure 2.**
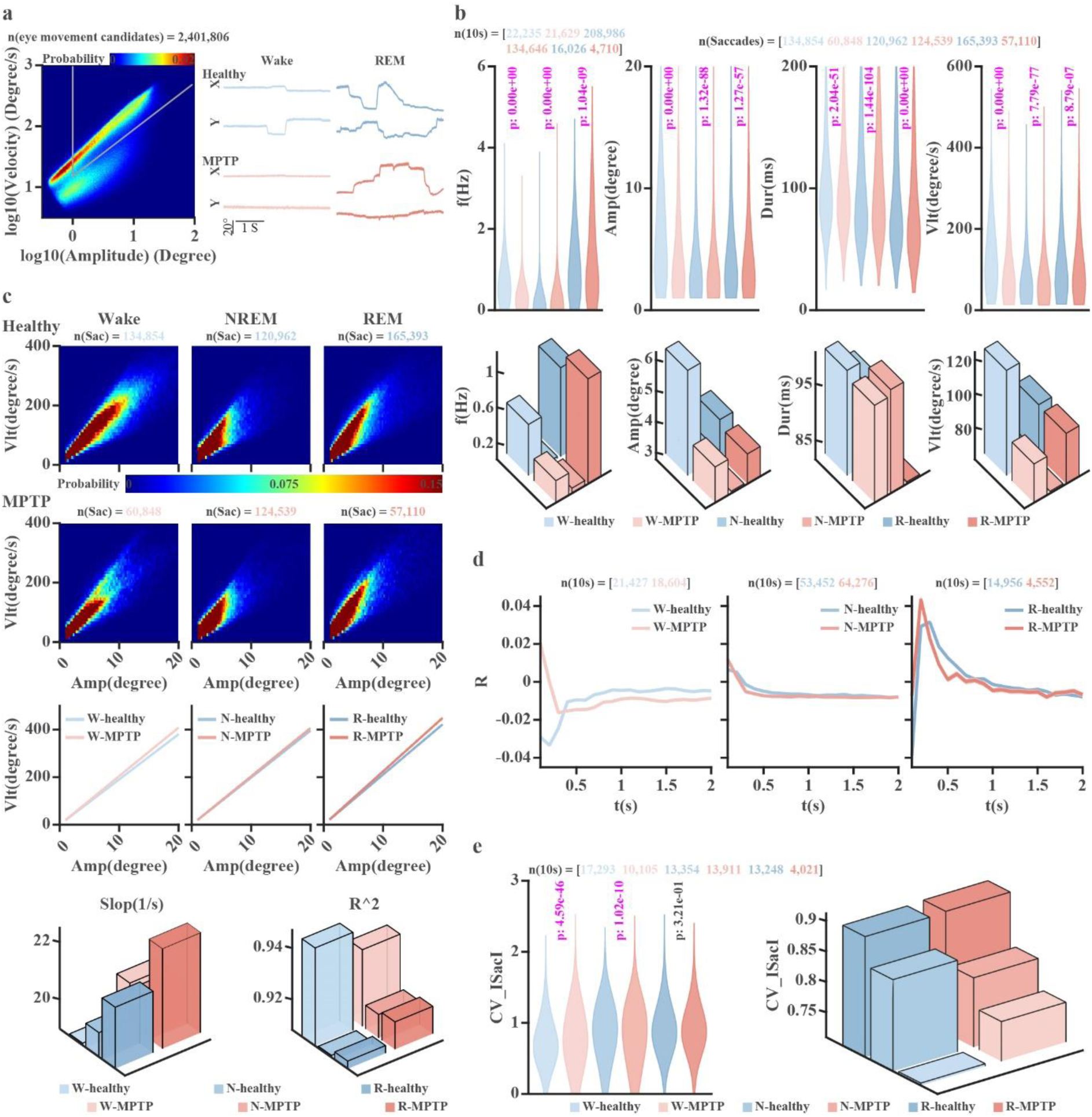
MPTP selectively impairs saccade dynamics during wakefulness while preserving them during REM sleep. **a.** Fast eye movements (saccades) are located to the right of the vertical gray line and above the diagonal gray line. Representative wakefulness- and REM sleep-associated eye movements before and after MPTP are shown in (**a**, right). **b**. Distributions of saccade frequency (f), amplitude (Amp), duration (Dur), and peak velocity (Vlt) across vigilance states before and after MPTP (**b**, top), with corresponding group mean values shown below (**b**, bottom). **c**. Amplitude peak–velocity relationships (main sequences) of saccadic eye movements across vigilance states, before and after MPTP, are shown in the first and second rows. Linear regression fits are presented in the third row, and the corresponding slopes and goodness-of-fit values (R²) are shown in the fourth row. **d.** Autocorrelation histograms of saccades across vigilance states before and after MPTP. R values are shown as mean ± SEM. **e.** The distribution (**e**, left) and the means (**e**, right) of the coefficient of variation of inter-saccade intervals (CV-ISacI) across vigilance states before and after MPTP. Before MPTP is in blue and after MPTP in red (from light to dark across wakefulness, NREM, and REM). n(10s) is the number of 10-second epochs. The statistical difference of variables between before and after MPTP in **b** and **e** is calculated using the Wilcoxon rank sum test (p<0.05). The p-values indicating significant differences are marked in magenta; otherwise, they are black. Abbreviations and color coding as in Figure 1.

**Figure 3.**
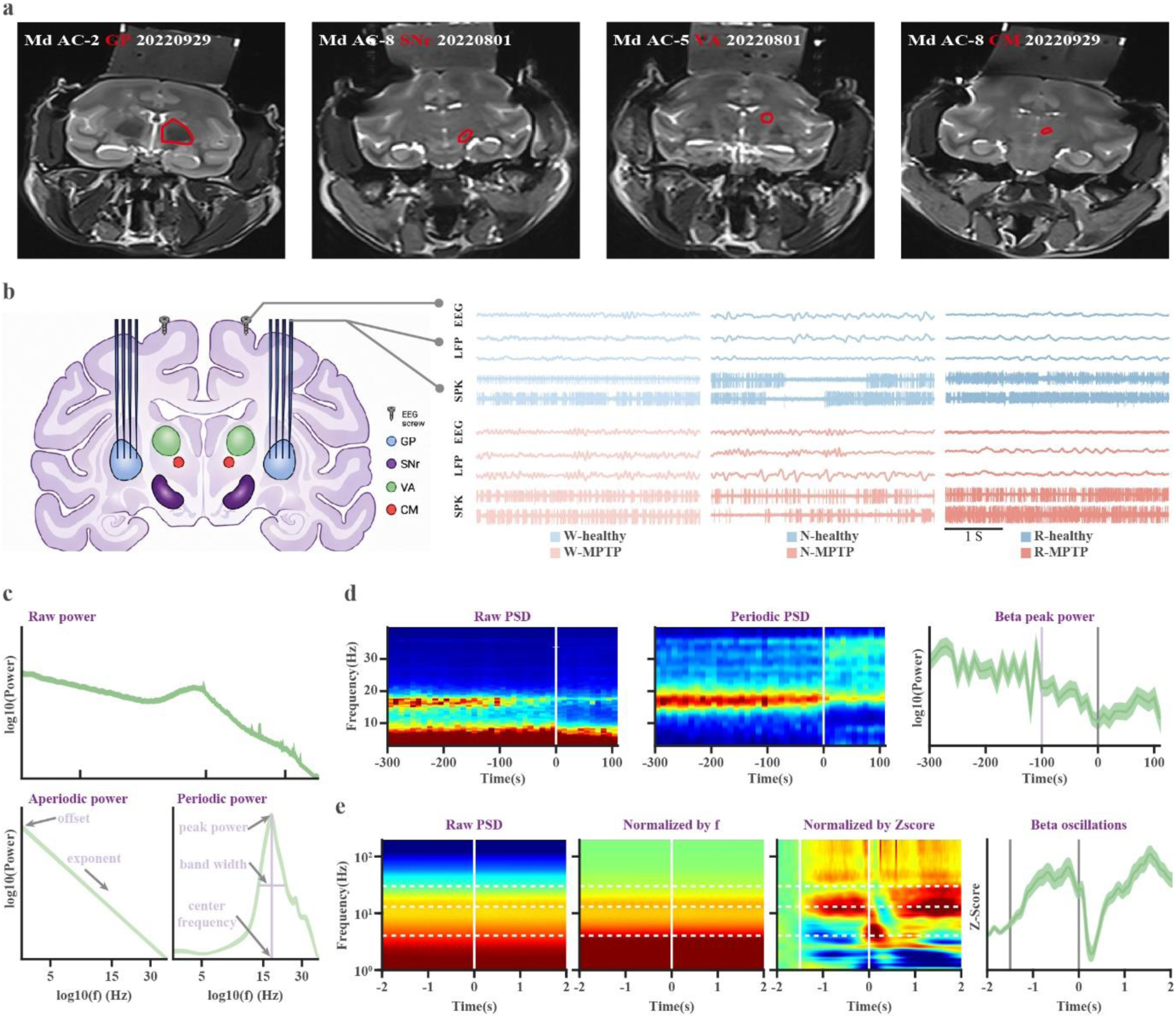
Experimental and analytical framework for investigating sleep–wake neural dynamics. **a**. MRI displays of the stereotaxic location of the recorded nuclei (GPe, globus pallidus-external; SNr, substantia nigra reticularis; VA, ventral-anterior of the thalamus; CM, centromedian of the thalamus). The recorded nuclei and their distances from the anterior commissure (AC) are marked in the corresponding images. **b.** Schematic diagram of neuronal recording techniques (left; ChatGPT-generated image). Examples of EEG, GPe LFP, and GPe SPK activity across vigilance states before and after MPTP (right). **c.** FOOOF (fitting oscillations & one over frequency) analysis separating raw signals (top) into aperiodic (bottom-left) and periodic (bottom-right) components and the quantification of the periodic and aperiodic components. **d**. Aligning spectral power to NREM-to-REM transitions. Raw (left), periodic (middle) power spectrogram densities (PSD), and their corresponding periodic beta power (right) aligned to the transition from NREM to REM sleep. The vertical white and grey lines indicate the sleep transition points (time 0). The vertical light purple line indicates the change point of the beta power. The blue-to-red color of the PSD indicates the percentage of power in a frequency relative to total power. **e.** Aligning spectral power to eye movement (saccades). From left to right: raw, normalized by frequency (total power across all frequency bins of a time unit), normalized by z-score (of the 2-1.5s pre-saccade period), and the beta power as a function of time around the saccade. In the first three subplots, the blue-to-red color scale indicates log10(power), the percentage of a frequency power relative to the total power, and Z-score-normalized power, respectively. Time 0 indicates the timing of a saccade and is marked by vertical white lines in the first three subplots and grey vertical lines in the last subplot. The baseline for z-score normalization is the area before the first white and grey vertical lines in the last two right subplots. The three horizontal dashed white lines indicate the 4 Hz, 13 Hz, and 30 Hz frequencies from bottom to top, respectively. Abbreviations and color coding as in Figure 1.

**Table 1.** The recording database.

| **Table 1. The recording database** |  |  |  |  |  |  |
| --- | --- | --- | --- | --- | --- | --- |
| State | Recording sessions (night/electrode) |  |  |  |  |  |
|  | Structure | Basal ganglia |  | Thalamus |  | Total |
|  |  | GPe | SNr | CM | VA |  |
| Healthy (before MPTP) | Md | 14/85 | 26/145 | 11/57 | 8/27 | 59/314 |
|  | Wh | 11/60 | 18/89 | 9/58 | 9/46 | 47/253 |
|  | Md+Wh | 25/145 | 44/234 | 20/115 | 17/73 | 106/567 |
| MPTP-I | Md | 3/17 | 3/11 | 3/13 | 4/15 | 13/56 |
|  | Wh | 2/14 | 3/18 | 2/9 | 2/15 | 9/56 |
|  | Md+Wh | 5/31 | 6/29 | 5/22 | 6/30 | 22/112 |
| MPTP-II | Md | 5/31 | 6/36 | 6/33 | 8/32 | 25/132 |
|  | Wh | 5/35 | 10/53 | 5/35 | 5/29 | 25/152 |
|  | Md+Wh | 10/66 | 16/89 | 11/68 | 13/61 | 50/284 |

*This recording database shows the number of recording sessions (nights) in the basal ganglia (globus pallidus external segment (GPe) and substantia nigra reticulata (SNr)) and thalamus (ventral anterior (VA) and centromedian (CM)) in healthy (before MPTP), MPTP-I, and MPTP-II states of the two non-human primates (Md and Wh). EEG was recorded at each recording session. The number of EEG recording days equals the sum of the recording nights (total = 106, 22, 50 for Healthy, MPTP-I, and MPTP-II states, respectively). The number of EEG signals is double the EEG recording days (total = 212, 44, 100 for Healthy, MPTP-I, and MPTP-II states, respectively).*

### MPTP effects on REM sleep and EMG

MPTP induces severe, stable parkinsonism during wakefulness (Fig. 1a). Figure 1b presents representative sleep hypnograms recorded before MPTP, during the first week post-MPTP, and throughout the final two weeks of the recording period. Figure 1c illustrates the nightly proportions of wakefulness, NREM sleep, and REM sleep (calculated from sleep onset) for each NHP, both before and after MPTP. REM sleep is almost abolished for 9-16 days following MPTP administration (MPTP-I phase), followed by a partial re-emergence of REM episodes during a later stage of parkinsonism (MPTP-II phase, Fig. 1b, c). Hereafter, the term “MPTP” refers specifically to the MPTP-II phase.

Unlike the robust REM rebound observed after REM sleep deprivation(*11*, *12*), REM recovery in the MPTP-II state is incomplete and fragmented. REM episode frequency is reduced relative to the pre-MPTP (healthy) condition, although this effect does not reach statistical significance (Figs. 1d and S1a). In parallel, REM episode duration is markedly decreased (Figs. 1e and S1b), and together with the decrease in the REM frequency, yielded a significant decrease in REM probability (Fig. 1c). Additionally, the latency to the first REM episode is significantly prolonged (Fig. 1f). Before MPTP, the final REM episode of the night is substantially longer than the first; this within-night escalation is blunted after MPTP (Figs. 1g and S1c). Finally, MPTP increases EMG activity across vigilance states, including REM sleep (Fig. 1h). Collectively, these findings are consistent with impaired REM architecture and loss of muscle atonia in RBD.

### MPTP effects on saccadic eye movement

Because rapid eye movements provide quantifiable motor behavior that occurs across vigilance states(*13*), we next examined whether MPTP-induced parkinsonism affects saccade generation similarly during wakefulness, NREM sleep, and REM sleep. Saccades (rapid eye movements) were identified based on their characteristic amplitude–velocity relationship and analyzed across all vigilance states (Fig. 2a).

MPTP produces a marked state-dependent alteration in saccade dynamics. During wakefulness, parkinsonism significantly reduced saccade frequency, amplitude, and peak velocity, accompanied by a shortened saccade duration (Fig. 2b), which indicates pronounced akinesia and bradykinesia. In contrast, REM-sleep-associated saccades remained largely preserved. Although minor quantitative changes are detectable, the overall distributions of REM saccade amplitude, duration, and velocity are substantially less impaired than those observed during wakefulness (Fig. 2b). NREM sleep’s saccades showed an intermediate phenotype.

To determine whether parkinsonism altered the fundamental kinematic organization of eye movements, we examined their amplitude–velocity relationship (main sequence). Before MPTP, saccades across vigilance states exhibit the expected strong positive amplitude–velocity relationship (Fig. 2c and S1d). Following MPTP administration, the main-sequence relationship closely resembles the healthy state, with only modest changes in slope and goodness-of-fit (R²) values (Fig. 2c and S1d). Since the features of the amplitude-velocity relationships are maintained at the brainstem level(*14*), the preservation of main-sequence kinematics suggests that the reduced saccade frequency and amplitude are generated at higher levels than the brainstem (e.g., basal ganglia and cortex).

Finally, we investigated the temporal organization of saccadic behavior. Autocorrelation analyses reveal substantial changes in the saccadic temporal organization during wakefulness after MPTP, whereas REM sleep autocorrelation profiles are comparatively preserved (Fig. 2d and S1e). Consistent with this observation, the coefficient of variation of inter-saccade intervals after MPTP increases significantly during wakefulness but shows little or no alteration during REM sleep (Fig. 2e), indicating that the temporal patterning of REM-associated eye movements remains relatively stable.

Together, these findings demonstrate a striking state dependence of parkinsonian motor dysfunction. Whereas wakefulness is characterized by reduced saccade vigor and disrupted temporal organization, REM sleep largely preserves both the kinematic and temporal properties of saccadic movements. This dissociation mirrors clinical observations that PD-RBD patients can generate vigorous, complex, well-coordinated limb movements and clear articulated speech during REM sleep(*8*). We therefore investigated the neuronal correlates of the state-dependent expression of parkinsonian saccadic deficits (Fig. 3).

### Spontaneous beta oscillations (13-30 Hz) in the basal ganglia and thalamus are significantly reduced during REM sleep

We simultaneously recorded frontal EEG together with neuronal activity from one of the four subcortical nuclei (GPe, SNr, VA, or CM) before and after MPTP (Fig. 3a, b). Up to 8 microelectrodes were inserted bilaterally into BG or thalamic targets during each recording session (Fig. 3a, b-left). Here, we analyzed EEG, LFPs, and SPKs during stable recording epochs. Recording yields are summarized in Table 1.

Because pathological oscillatory activity can be superimposed on broadband aperiodic neural activity, we separated periodic oscillations from the aperiodic background using FOOOF analysis(*15*, *16*). This allows independent quantification of oscillatory parameters (peak power, center frequency, and aggregate periodic power), as well as aperiodic parameters (offset and exponent) (Fig. 3c). To further examine the neural dynamics underlying behavioral state transitions, spectral activity was aligned to NREM-to-REM transitions and to individual saccades (Fig. 3d, e).

The raw spectrograms (normalized by the total power in each time bin) are shown in Figure 4a. Separating periodic and aperiodic components reveals that MPTP significantly changes the aperiodic parameters of frontal EEG and LFP signals throughout the basal ganglia-thalamocortical network (Figs. 4b, c and S2a). These changes are observed during wakefulness, NREM sleep, and REM sleep, although their magnitudes vary across vigilance states and recording sites. In subcortical recordings, MPTP increases wakeful LFP offsets without altering exponents, whereas both offsets and exponents are reduced during NREM and REM sleep (Figs. 4b, c and S3). In contrast, the aperiodic parameters of spiking activity are largely preserved after MPTP, with only modest changes across nuclei (Figs. 4b, c and S2a). Thus, parkinsonism predominantly alters the aperiodic broadband components of population field potential rather than those of neuronal spiking activity. However, MPTP does not alter the effects of the vigilance states; aperiodic parameters are higher during NREM sleep than during activated states (wakefulness and REM sleep) (Fig. 4).

**Figure 4.**
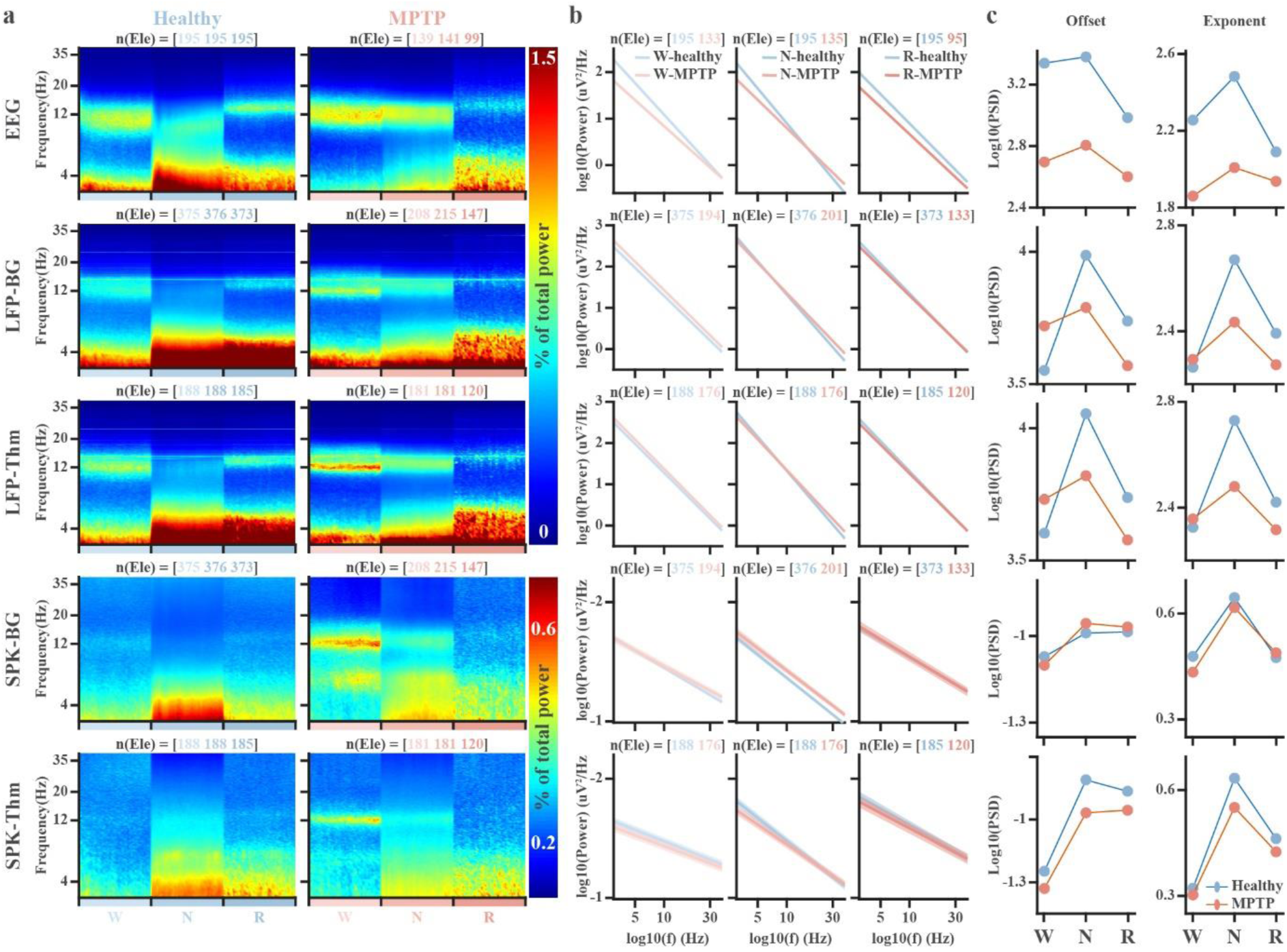
MPTP selectively alters aperiodic offset and exponent in EEG and LFP, but not spiking activity. **a**. Raw spectrograms of frontal EEG, basal ganglia (BG) and thalamic (Thm) LFP and SPK activity before and after MPTP. Sleep-stage time is normalized into 40 bins by averaging power values within each time bin. Pre- and post-MPTP periods are indicated by blue and red color schemes, respectively, with different shades representing wakefulness, NREM sleep, and REM sleep (W, N, and R). **b.** Average aperiodic power as a function of frequency for frontal EEG, BG, and Thm LFP and SPK activity across vigilance states. The aperiodic power is shown as mean ± SEM. Note the different y-axis ranges among EEG, LFP, and SPK activity. **c.** The average aperiodic parameters (offset and exponent) before and after MPTP across vigilance states. Their distribution and statistical results are shown in supplementary Figure S2a. The number of electrodes (Ele) is the same as those shown in **b**. Abbreviations and color coding are as in Figure 1.

In the healthy state, beta-band periodic activity in cortical EEG and subcortical LFPs exhibits strong vigilance-state dependence, peaking during wakefulness, decreasing during NREM sleep, and partially recovering during REM sleep (Figs. 5a–c and S2b). In contrast, beta rhythmicity is largely absent in BG and thalamic spiking activity across all vigilance states (Figs. 5a–c and S2b). Unlike the aperiodic parameters, MPTP markedly alters the vigilance state-dependent trend in beta-band periodic power. Consistent with previous studies(*1*, *17*), MPTP markedly increases beta-band peak power throughout the frontal cortex, BG and thalamic LFPs during wakefulness, with moderate increases during NREM sleep (Figs. 5a–c, S2b and S4). Remarkably, this parkinsonian beta enhancement disappears during REM sleep, where beta peak power remains comparable to the healthy state (Figs. 5a–c and S2b). A similar state-dependent pattern is observed in the spiking activity; MPTP enhances beta rhythmicity during wakefulness but produces little or no increase during REM sleep. Finally, MPTP also induces modest state-dependent shifts in beta center frequency (Fig. 5c and S2b).

**Figure 5.**
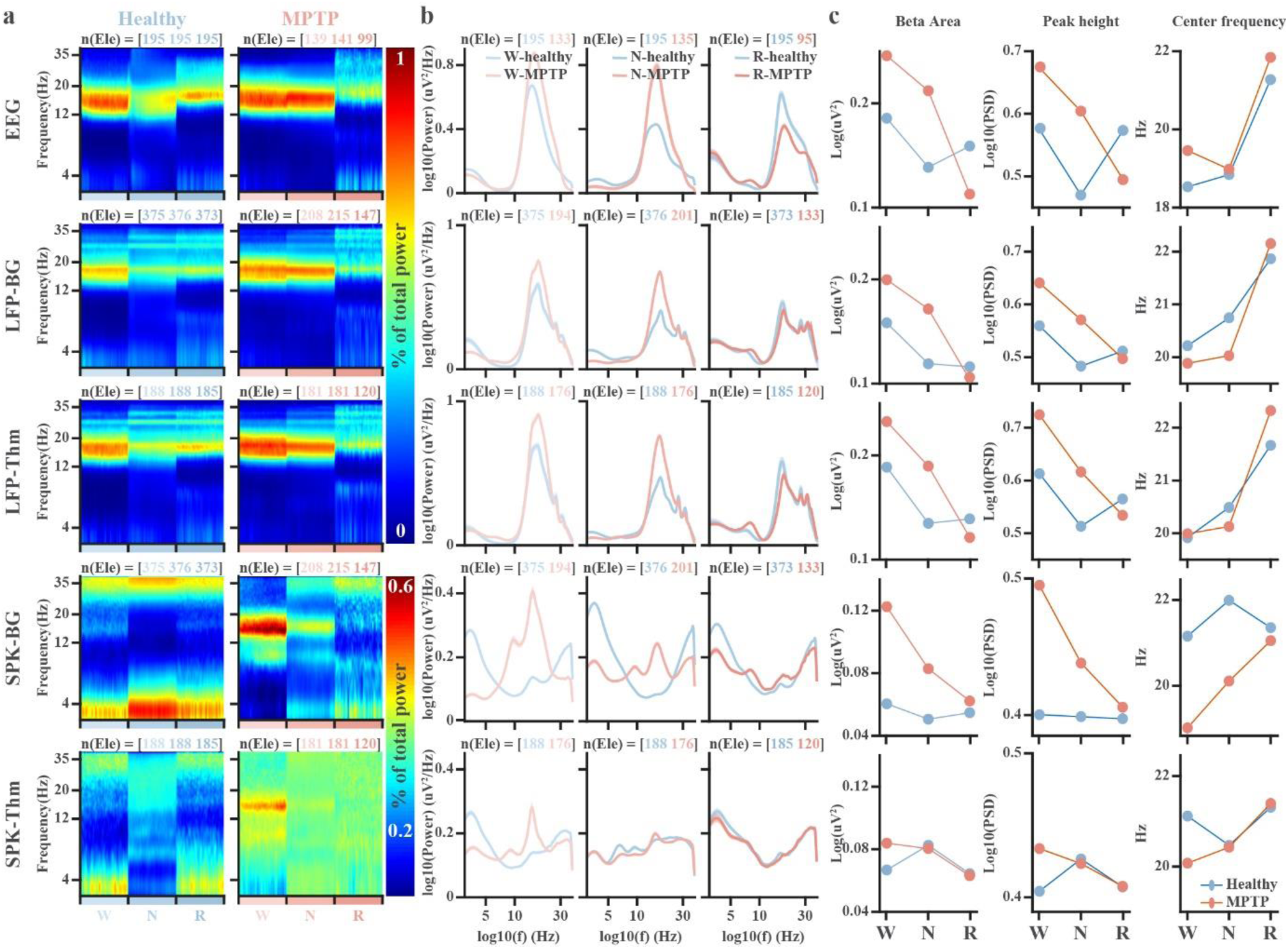
MPTP enhances beta-band periodic power during wakefulness and NREM, but not during REM sleep. **a**. Periodic spectrograms of frontal EEG, basal ganglia (BG), and thalamic (Thm) LFP and SPK activity before and after MPTP. Sleep-stage time is normalized into 40 bins by averaging power values within each time bin. Blue and red denote the pre- and post-MPTP conditions, respectively. Within each condition, light, medium, and dark shades indicate wakefulness (W), NREM sleep (N), and REM sleep (R), respectively. **b.** Average periodic power as a function of frequency of frontal EEG, BG and Thm LFP and SPK activity across vigilance states. The periodic power is shown as mean ± SEM. Note the different y-axis ranges among EEG, LFP, and SPK activity. **c.** The average periodic beta area (column 1), peak power (column 2), and beta center frequency (column 3), before and after MPTP across vigilance states. The beta area is the summation of power in the beta frequency range (the summation of log10(beta power) * log10(frequency)). Their distribution and statistical results are shown in supplementary Figure S2b. The number of electrodes (Ele) is the same as those shown in **b**. Abbreviations and color coding are as in Figure 1.

### Beta suppression precedes behavioral manifestations of REM sleep after MPTP

To determine the temporal relationship between beta oscillations and vigilance-state transitions, neural activities were aligned to NREM-to-REM transitions (Figs. 6 and S5). In healthy NHPs, beta-band activity begins to change approximately 30 s before REM onset across cortical EEG, BG, and thalamic recordings (Fig. 6a–c). Following MPTP administration, these changes emerge earlier, approximately 100 s before REM onset, indicating a prolonged transition period in the parkinsonian state. Beta power remains elevated throughout NREM sleep after MPTP (Fig. 5) before progressively declining before the onset of the next REM episode (Fig. 6).

**Figure 6.**
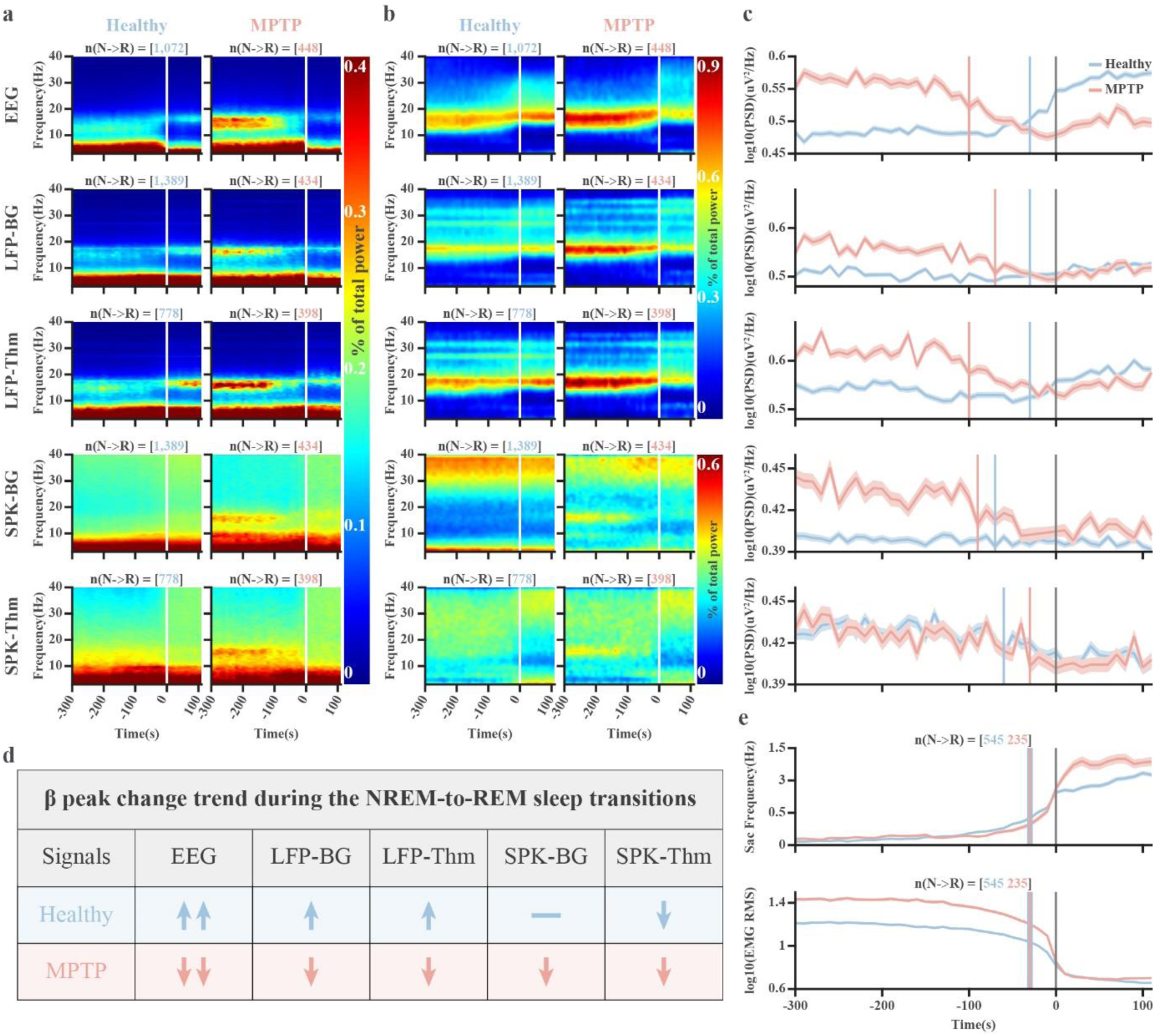
Beta power modulation precedes NREM-to-REM transitions before and after MPTP. **a**. The healthy (left panel) and MPTP (right panel) raw power spectrogram densities (PSD) of frontal EEG, and basal ganglia (BG) and thalamic (Thm) LFP and SPK activities are aligned to the transition from NREM to REM sleep (time = 0). **b.** The healthy (left panel) and MPTP (right panel) periodic PSDs are aligned to the transition from NREM to REM sleep. **c**. Beta peak power aligned to the NREM-REM sleep transition. **d.** Modulation of beta peak power around NREM-to-REM sleep transitions is shown using arrows. Upward/downward arrows indicate increase/decrease of beta power. **e.** Saccade frequency and EMG RMS aligned to NREM--REM sleep transitions. The vertical white (**a**, **b**) and grey (**c**, **e**) lines (time 0) indicate the first epoch of REM sleep. There are 30 NREM segments and 12 REM 10-second epochs before and after time 0. The change points in the beta power before and after MPTP are detected and marked by vertical blue and red lines (**c**, **e**), respectively. The vertical blue lines in **e** are wider than the vertical red lines to enhance visibility, since they are at the same time. The number of NREM-to-REM sleep transitions in **a, b,** and **e** is shown in the title of the corresponding subplots. The number of these sleep transitions in **c** is the same as that in **b.** The data in **c** and **e** are shown as mean ± SEM. Abbreviations and color coding as in Figure 1.

Although the temporal profiles differ across recording modalities and frequency ranges, the overall sequence of neural events is remarkably consistent. Cortical EEG exhibits a pronounced downshift of dominant delta-theta activity during the transition to REM sleep both before and after MPTP, whereas comparable shifts are not observed in subcortical LFPs or spiking activity (Fig. 6a). At the same time, beta dynamics display distinct state- and modality-dependent patterns. Cortical beta power increases before the NREM-to-REM transition in healthy animals but progressively declines after MPTP (Fig. 6b-d). Subcortical LFPs exhibited qualitatively similar dynamics, although the pre-transition modulation is less pronounced in health (Fig. 6b-d). However, BG spiking shows no discernible beta oscillation before MPTP, with relatively stable power across the beta frequency range, whereas the pathological beta oscillation emerging after MPTP exhibits a marked reduction in oscillatory power before REM onset. Thalamic spiking exhibits only modest reductions under both healthy and MPTP conditions (Fig. 6b-d).

Change-point analysis reveals that beta modulation consistently precedes the onset of REM sleep across recording modalities. After MPTP, beta reduction also precedes REM-associated behavioral changes, including increased saccade frequency and reduced muscle tone (Fig. 6e). Before MPTP, neural and behavioral transitions occur nearly simultaneously, whereas after MPTP, they are separated: behavioral changes are preserved, but the corresponding neural dynamics are prolonged preceding REM sleep.

### Saccade-related beta desynchronization is attenuated in the basal ganglia and thalamus during REM sleep

Having established that beta suppression preceded the transition into REM sleep (Fig. 6), we next examined whether movement-related oscillatory modulation was preserved during REM sleep. We computed saccade-related time–frequency spectrograms of EEG, LFPs, and spiking activity across wakefulness, NREM Sleep, and REM sleep before and after MPTP (Figs. 7, S6, and S7).

**Figure 7.**
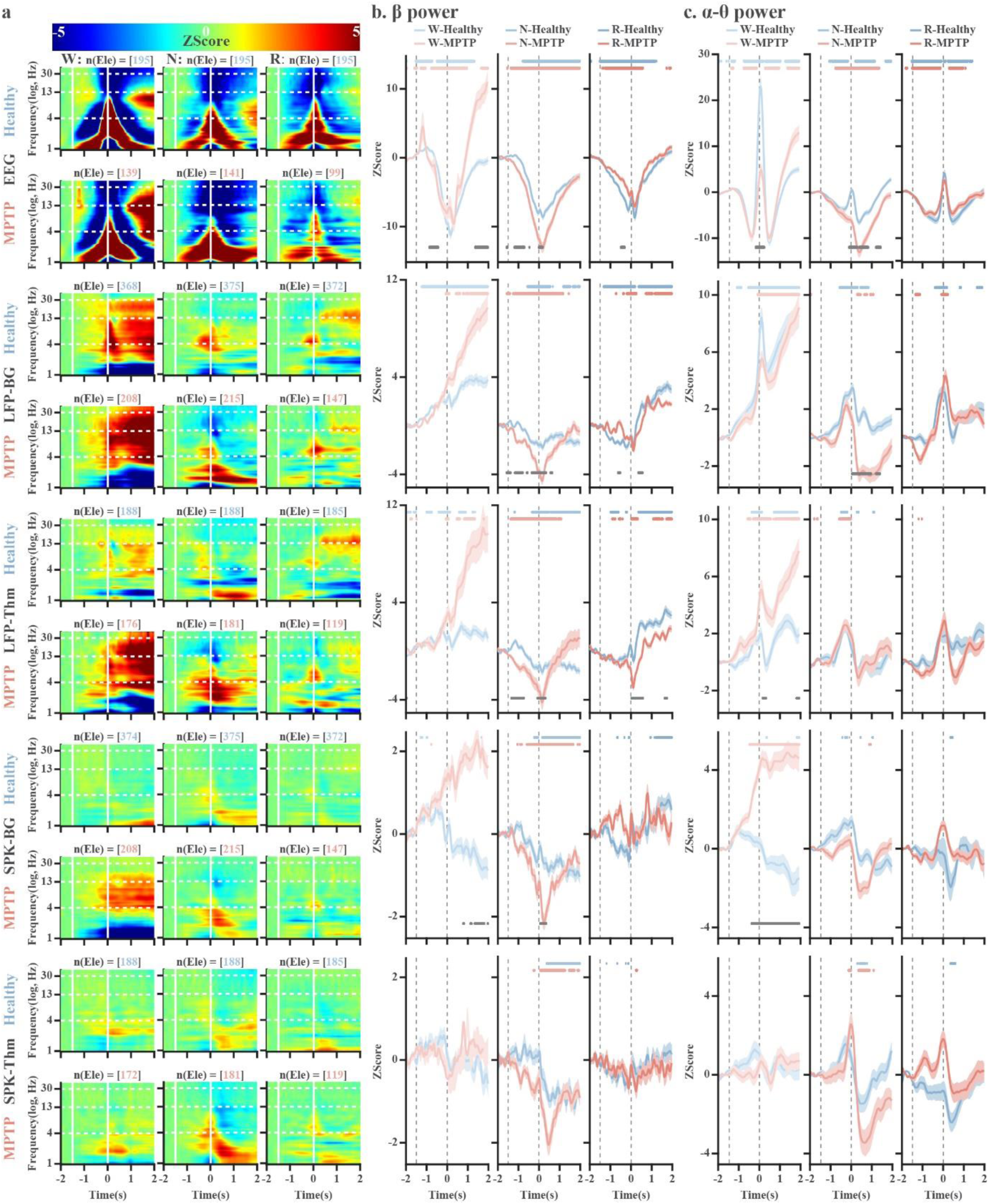
State-dependent effects of MPTP on saccade-aligned beta desynchronization and theta-alpha synchronization. **a**. The spectrograms of frontal EEG, basal ganglia (BG), and thalamic (Thm) LFP and SPK are aligned to fast eye movements (saccades). The 1st vertical solid white line indicates the baseline (−2 to −1.5 seconds before saccades) used for the z-score normalization. The 2nd vertical solid white line indicates the timing of saccades (time 0). The beta frequency range is between the 1st and 2nd upper horizontal dashed white lines. The theta-alpha frequency range is between the 2nd and 3rd horizontal dashed white lines. **b.** The average power at the beta (13-30Hz) frequency range as a function of time around saccades. **c.** The average power in the theta-alpha (4-12Hz) frequency range. The data in **b** and **c** are shown as mean ± SEM. The vertical dashed grey lines in **b** and **c** indicate the same meaning as the vertical solid white lines in **a**. The blue and pink dots at the top of each subplot indicate a significant difference from the baseline (from −2 to −1.5 seconds before the saccade). The Bonferroni-corrected Wilcoxon rank-sum test was used to assess differences in beta or theta-alpha power across different lag time points (p <0.05/1375) in **b** and **c**. The grey dots at the bottom of those sub-plots indicate significant differences in beta or theta-alpha power before and after MPTP (Wilcoxon rank-sum test, p < 0.05). Abbreviations and color coding as in Figure 1.

During memory-guided saccades in the healthy state, EEG and LFPs exhibit the expected event-related suppression/augmentation (desynchronization/synchronization) pattern in the beta and theta–alpha bands, consistent with previous studies(*18*, *19*). Spiking activity exhibits a comparable beta-desynchronization profile, with an earlier onset than EEG and subcortical LFP signals (Fig. S6).

We further examined spontaneous saccades across vigilance states (Figs. 7 and S7). Before MPTP, saccade-related beta desynchronization is prominent in cortical EEG but weak in subcortical LFPs and spiking activity. MPTP produces striking state-dependent alterations (Fig. 7a, b). During wakefulness, beta desynchronization is modestly reduced, whereas post-saccadic beta rebound increases. In contrast, during NREM sleep, beta desynchronization is enhanced, but beta rebound is largely absent. This can probably explain the NREM fast eye movement in Figure 2, despite the elevated baseline beta activity observed in Figure 5. Remarkably, REM sleep is largely unaffected: saccade-related beta desynchronization in cortical EEG remains similar before and after MPTP, and saccade-related modulation of subcortical LFPs and spiking activity is minimal under both conditions.

Theta–alpha synchronization also shows state-dependent changes (Fig. 7a, c). During wakefulness, MPTP reduces theta–alpha (4–12 Hz) power augmentation (synchronization) in EEG and BG LFPs. BG spiking activity after MPTP starts to increase and reaches its highest amplitude when eyes start to move. This high power is maintained up to the end of the eye movement. Awake thalamic spiking activity is not obviously modulated by saccades in either condition. Finally, theta–alpha modulation of EEG and subcortical activity is generally weaker and usually does not differ significantly between the healthy and MPTP conditions during either NREM or REM sleep.

## Discussion

Our results align with the hypothesis that EEG and subcortical beta oscillations are a stop or status quo signal(*20*, *21*). Although beta activity is pathologically enhanced during wakefulness and NREM sleep in parkinsonism, this enhancement disappears during REM sleep. This state-dependent suppression of beta activity may release motor circuits from inhibitory control, permitting rapid eye movements during REM in both healthy and parkinsonian conditions. Consistent with this interpretation, REM-related saccades after MPTP remain as frequent and rapid as those in the healthy condition, paralleling the fast, smooth limb movements and clear articulation of speech reported in PD-RBD patients(*8*). More broadly, α-synuclein pathology in the brainstem likely underlies the loss of REM atonia in PD-RBD(*4*), whereas the concomitant reduction in beta activity in the BG and thalamus may account for the relative preservation, and even facilitation of movement kinematics during REM sleep in PD-RBD patients.

### Abnormal REM sleep and saccadic eye movements feature in PD and the MPTP model

Sleep disorders, especially insomnia, fragmentation of sleep, and excessive daytime sleepiness, were repeatedly reported in NHPs administered with MPTP(*9*, *22–25*). The effect of MPTP on REM sleep was occasionally described. However, consistent with our current study (Fig. 1c), several reports indicate that MPTP induced a significant reduction in REM sleep, followed by RBD-like symptoms(*26*, *27*). Consistent with our findings in the MPTP I and II phases, without and with re-emergence of REM sleep, few studies have described MPTP induction of an abrupt ablation of REM sleep followed by a subsequent partial re-emergence of REM sleep(*28*, *29*). Similar findings of MPTP-induced REM elimination were reported in cats treated with a single dose of MPTP, without PD symptoms or dopamine depletion(*30*).

MPTP is not the only agent that induces abrupt reduction and late re-emergence of REM sleep. Most classes of antidepressants also produce a marked suppression of REM sleep, often followed by rebound or partial re-emergence during chronic administration or after withdrawal(*31–34*). With prolonged administration, REM sleep often partially returns despite continued medication, suggesting that the REM-generating circuitry adapts to sustained monoaminergic manipulations. Together, these pharmacological observations demonstrate that abrupt REM suppression followed by delayed re-emergence is a general phenomenon of monoaminergic therapy, not unique to MPTP-induced Parkinsonism.

Polysomnography studies of PD patients show reduced REM sleep, prolonged REM latency, and altered REM architecture(*35*, *36*). Our results in the MPTP stage (after the reemergence of REM sleep) are consistent with those previous reports. However, reported changes in REM density (the frequency of rapid eye movements during REM sleep) in PD have been inconsistent. While some studies have found increased REM density in patients with PD-RBD(*36*), more recent work suggests that REM density is generally reduced and correlates with the severity of motor impairment(*35*). Our results demonstrate increased REM density in MPTP NHPs, enabling the exploration of RBD mechanisms.

The data on the kinematics of PD REM saccades are much sparser. Healthy human EOG studies found weaker amplitude-velocity coupling of saccades during REM sleep than during wakefulness(*37*). In PD, wake-spontaneous saccades are more fragmented (multistep) with decreased amplitude. Visually guided saccades are also hypometric and have longer latencies than in the healthy state(*38*). NHP MPTP models faithfully reproduce the human PD awake pattern of saccadic bradykinetic abnormalities(*39*). Here, we showed that following MPTP administration, spontaneous saccades during wakefulness are akinetic (reduced frequency) and bradykinetic (slower peak velocity and smaller amplitude). Notably, when we compare REM eye movements recorded before and after MPTP, the MPTP REM saccades are neither akinetic nor bradykinetic, as reported for the limb movements and speech of PD-RBD patients(*8*).

### Cortical and subcortical beta activity across vigilance states before and after MPTP

Pathological β oscillations in the subthalamic nucleus (STN) of patients with PD exhibit marked sleep– wake state dependence(*17*). Compared with wakefulness, Cortical EEG, STN and GPi LFP β activity significantly decreased during NREM sleep. In contrast, β activity re-emerged during REM sleep, with power levels returning to values comparable to wakefulness and, in some patients, even exceeding waking levels(*40–42*). These clinical findings resemble the EEG/LFP β dynamics observed in our monkeys under healthy conditions and in EEG studies of healthy controls(*43*), where β activity decreases during NREM sleep and re-emerges during REM sleep.

The classically activated brain states (wakefulness and REM sleep) vs. NREM sleep patterns found in EEG and subcortical LFP studies differ markedly from the sleep-state-dependent dynamics observed in spiking activity of the healthy BG and thalamus, suggesting that EEG and subcortical LFP β oscillations and subcortical neuronal spiking β oscillations may reflect distinct physiological processes(*22*). Following MPTP administration, EEG/LFP β activity in our monkeys exhibited a monotonic decline across vigilance states, with the highest β power during wakefulness, intermediate levels during NREM sleep, and the lowest levels during REM sleep. This pattern closely matched the post-MPTP vigilance state-dependent changes observed in BG spiking β oscillations.

One possible interpretation is that our MPTP monkeys represent a more advanced Parkinsonian state (Hoehn and Yahr stage 5)(*44*), whereas patients undergoing DBS surgery may occupy an intermediate stage between healthy and advanced disease conditions (Hoehn and Yahr stage 2-3). Under this framework, the observed sleep-state-dependent trajectory of EEG/LFP β activity may reflect the relative contributions of physiological and pathological β oscillations. As disease severity increases, pathological β activity may progressively dominate network dynamics, causing LFP/EEG β patterns to shift from the healthy sleep-dependent profile toward a pathological profile resembling the subcortical spiking β activity. Methodological differences across studies may further contribute to the variability of reported findings. These include whether recordings are obtained after a short-term withdrawal of dopaminergic medication (e.g., three hours) or following a withdrawal beginning in the morning before night sleep. Our monkeys received only a morning dose of short-acting dopamine replacement therapy before a night recording. Other differences include EEG contact configuration; monopolar vs. differential bipolar LFP recording; different methods of spectral analysis and display of results (absolute power, fraction of total power, range of frequencies used to calculate the total power, with/without whitening).

If this hypothesis of disease severity is correct, the reliability of LFP β power as a biomarker for distinguishing wakefulness and sleep stages may progressively decline with disease progression. Our results demonstrate that the sleep-state-dependent trends in the LFP aperiodic exponent and offset remain largely preserved after MPTP. The exponent exhibited greater stability than the offset across disease states (Fig. 4). These findings suggest that LFP aperiodic activity, particularly the exponent, may constitute a more robust biomarker for sleep–wake state identification than LFP β-band power alone. Looking forward, advances in implantable neurotechnology may eventually enable chronic recording of neuronal spiking activity from BG circuits in clinical settings. Given that spiking activity represents the output of the recorded structure(*16*), and that spiking β oscillations in our study exhibited more consistent state-dependent dynamics across physiological and pathological conditions, spiking-based biomarkers may ultimately provide a more reliable substrate for adaptive neuromodulation than LFP-derived β power.

The distinct β-dynamics of BG spiking activity may lead to fundamentally different therapeutic strategies for adaptive DBS (aDBS). Both previous clinical studies and our findings suggest that β activity should be suppressed during wakefulness and NREM sleep in PD, although the optimal stimulation intensity and threshold for adaptive treatment may differ between these states. However, the implications for REM sleep are substantially different. According to previous clinical observations, LFP β activity in the BG re-emerges or even increases during REM sleep, and this increase appears to be more pronounced in patients with RBD and during episodes of RSWA(*40*, *45*). Such trends would support maintaining or even increasing therapeutic stimulation during REM sleep, as during wakefulness. In contrast, our findings suggest a different interpretation. In both healthy and parkinsonian states, β oscillations within BG are naturally suppressed during REM sleep. Thus, REM sleep may represent a physiological state in which the β-driven adaptive stimulation might be disadvantageous. For patients with RBD, it is conceivable that restoring physiological REM-related network dynamics might require facilitation, rather than suppression of specific oscillatory activities, e.g., in the gamma domain. If deep brain stimulation is delivered during REM sleep, its impact on the physiological frontal cortical beta activity (Fig. 6) remains unclear. Given that this beta activity may support cognitive functions or movement suppression, indiscriminate stimulation during REM could potentially interfere with these physiological processes. This interpretation aligns with previous studies(*46*, *47*) demonstrating that dopamine/STN-DBS treatment increases or does not affect the likelihood of RBD in PD patients. Whether next-generation state-dependent aDBS systems should target the BG, the cortex, or their interaction remains unknown and requires direct experimental testing. Our previous study(*48*) suggested that the BG do not drive their thalamic and cortical targets and that there is state-dependent dissociation between the extrapyramidal (BG, brainstem) and pyramidal (cortex) systems. These findings suggest that future DBS paradigms may need to consider not only oscillatory but also anatomical target, disease stage, and awake/sleep state, rather than applying a uniform β-suppression strategy across all behavioral conditions and anatomical targets.

### Why are the movements during REM sleep not as bradykinetic as those during wakefulness in PD?

In wakefulness, BG plays a key role in movement initiation via modulating automatic, axial, rhythmic activity. Excessive spontaneous beta-synchronized oscillations in BG nuclei are associated with akinesia/bradykinesia, and therapies that reduce beta oscillations (dopamine replacement, DBS) improve movement. Cortical (EEG) beta power decreases during movement(*49*). However, reports on movement-related changes in beta power in PD relative to control groups, or as a function of therapy, have been inconsistent(*50*). Some studies report reductions in beta modulation(*51*), while others report exaggerated beta responses(*52*). Several studies reported differences as a function of task details(*53*) or disease-related hemispheric dominance(*54*). Human neurophysiological studies focusing on RBD and PD-RBD describe similar changes in cortical EEG or BG LFP(*17*, *40–42*, *45*, *55–59*). Subcortical LFPs probably reflect cortical synaptic inputs and might be confounded by volume conduction of cortical activity. There are no studies of local spiking activity in the BG during REM sleep in both healthy controls and PD patients (or parkinsonian animals).

Here, we studied the cortical EEG, subcortical LFP, and subcortical SPK activity across vigilance states in both healthy and MPTP NHPs. For the memory-guided saccade task during wakefulness in healthy NHPs, the event-related decrease (desynchronization) in beta power and event-related increase in theta-alpha power (synchronization) in cortical EEG and subcortical LFP are similar to previous reports(*18*, *60*). However, our subcortical spiking activities demonstrate that event-related beta desynchronization reaches its lowest level earlier than cortical EEG and subcortical LFP (Fig. S6a, b), and beta synchronization was observed just before the event (saccade). Unlike cortical EEG and subcortical LFP, subcortical spiking activity does not have obvious event-related theta-alpha modulation (Fig. S6c).

Manzanilla et al.(*57*) found no differences in RBD limb movements-related EEG beta modulation regarding the presence or absence of PD. Other studies reported significantly elevated β activity in STN before and during movements in REM sleep, and reduced cortico-subthalamic coherence during REM and NREM movements(*42*). These results, together with the findings of BG beta activity levels similar to wakefulness and even higher during REM sleep(*17*, *42*), led to the hypothesis that RBD fast movements are enabled by bypassing the pathological movement-inhibiting BG networks in PD patients during sleep. Our results differ and suggest an active role for the BG throughout the whole natural cycle of vigilance states. We found reduced physiological beta oscillations in the cortex and no pathological beta oscillations in the BG during REM sleep (highest in wakefulness, medium in NREM sleep, lowest in REM sleep after MPTP, Fig. 5). Additionally, saccades-related EEG patterns show no differences between healthy and MPTP NHPs during REM sleep (Fig. 7a, b). After MPTP, subcortical LFP shows relatively weaker beta rebound during REM sleep. BG spiking activity during REM sleep is not modulated by MPTP. Our results may therefore explain the rapid movements of PD-RBD patients during REM sleep. The BG network might not be bypassed during the PD-RBD movements but be restored to a healthy activated state with minimal beta activity during REM sleep. The function of BG in PD patients is vigilance-state dependent and not purely dictated by the loss of dopamine.

### Study summary and limitations

Our findings demonstrate that pathological beta oscillations are strongly state-dependent. Although parkinsonism markedly enhances beta oscillations of BG spiking activity during wakefulness and NREM sleep, this pathological activity is largely absent during REM sleep, where movement-related beta desynchronization is also abolished. These findings suggest that REM sleep suppresses the beta-mediated “stop” signal, thereby permitting rapid movements despite severe parkinsonism. This state-dependent regulation of beta activity has important implications for understanding RBD and for the development of aDBS algorithms that account for sleep state(*61*).

This study has several limitations. First, although our findings were highly consistent across the two animals, the sample size was necessarily limited, as is common in NHP research and consistent with the 3Rs principles(*62*, *63*). Validation in larger cohorts and in human studies will therefore be important. Second, recordings were restricted to selected BG and thalamic nuclei (leaving the contribution of other brain regions unresolved) and to frontal EEG (that may be confounded by EOG and blinking). Third, although the MPTP model faithfully reproduces many motor features of PD, it does not recapitulate the α-synuclein pathology characteristic of most patients. Finally, REM sleep was identified using conventional polysomnographic criteria rather than clinical RSWA criteria. Future studies incorporating broader brain coverage, genetic PD models, and human recordings will be important for extending these findings.

## Materials and Methods

### Animal

Data were collected from two female NHPs (Chlorocebus sabaeus, Md and Wh), weighing 4-5 kg. This study was approved and supervised by the Institutional Animal Care and Use Committee of the Hebrew University and Hadassah Medical Center (MD-15-14412-5), and the veterinary staff of the Hebrew University’s primate facility. All experimental procedures were conducted in accordance with the National Institutes of Health Guide for the Care and Use of Laboratory Animals(*64*) and the ethical guidelines of the Hebrew University of Jerusalem.

### Experimental procedures – training, surgery, and MRI

To minimize stress, the NHPs were familiarized with a dark, double-walled, sound-attenuating experimental room before training on the eye-movement task. They were habituated to sitting and sleeping in an experimental primate chair in the experimental room with a gradual increase in time spent in this set-up. An eye-tracker (ISCAN, 21 Cabot Road, Woburn, MA 01801, USA) was used to record eye movements, and the NHPs were trained to perform a modified memory-guided saccade. A head holder, two ground screws, bilateral eye coils, two frontal skull-mounted EEG electrodes, and a bilateral frontoparietal 27 × 34 mm recording chamber were implanted in four separate surgeries. Those operations were performed under general anesthesia, with appropriate antibiotic and analgesic administration. To verify the precise location of the recording chamber, a 3T magnetic resonance imaging (MRI) scan was performed following recovery from the last surgery. More experimental details have been previously reported(*48*).

### Experimental procedures – recording protocol

The NHPs were transferred to the experimental room between 16:00 and 17:00. Task performance was typically completed within approximately two hours (This part was skipped after MPTP). Thereafter, the animals slept overnight (19:00-20:00 until 04:00-05:00; five nights per week) with the lights in the experimental room turned off, and under continuous infrared video monitoring and human supervision. During the daytime, the animals were maintained under a controlled feeding schedule and received their primary food intake during task performance. If the minimum daily caloric requirement was not achieved during task performance, supplemental nutrition was provided upon their return to the primate facility.

The NHPs were group-housed with conspecifics in an indoor yard during the daytime on weekdays and continuously housed together over the weekends. Environmental enrichment was systematically implemented by rotating enrichment devices weekly, including climbing nets, suspended ropes for jumping, and foraging substrates (e.g., hay containing hidden food items).

During recording sessions, eyelid state (open/closed) was monitored using an infrared video-based eye-tracking system (49-50 frames/s). Eye position was simultaneously measured using a scleral-coil system (Crist Instrument, Hagerstown, MD, USA) and recorded with SnR software (version 2.0.0; Alpha Omega Engineering). Electromyographic (EMG) activity from bilateral trapezius muscles and two frontal electroencephalographic (EEG) cranial screws were continuously acquired. Eye-coil, EMG, and EEG signals were sampled online at 2,750 Hz (SnR; Alpha Omega Engineering, Nof Hagalil, Israel).

MRI, primate brain atlases, and intraoperative electrophysiological mapping contributed to precisely localizing the nuclei of the BG (globus pallidus external segment (GPe) and substantia nigra reticulata (SNr))(*65*, *66*) and thalamus (ventral anterior (VA) and centromedian (CM))(*67*). During each nocturnal recording session, up to four independently controlled microelectrodes were advanced into the targeted nuclei (GPe, SNr, VA, or CM) in each hemisphere, yielding a maximum of eight electrodes per session. Recordings were restricted to homologous nuclei across right and left hemispheres (e.g., left and right VA). Our vertical approach to the subcortical structures makes the distinction between the external and internal segments of the globus pallidus (GPe and GPi) less reliable than the classical oblique approach (40-50 degrees from the midsagittal plane). Since our main analysis merges the BG recording sites, we believe that the potential mixing of GPi units with our GPe dataset does not affect our conclusions.

Two investigators (XL and JG) independently operated the Alpha Omega Electrode Positioning System (SnR; Alpha Omega Engineering, Nof Hagalil, Israel), each controlling four microelectrodes within a single hemisphere. We used commercial glass-coated tungsten microelectrodes with an impedance of 0.5-0.75 MΩ at 1 kHz (Alpha Omega Engineering, Nof Hagalil, Israel).

Raw neural signals were filtered online using a hardware filter (0.075–9,000 Hz, four-pole Butterworth) and digitized at 44 kHz. For real-time visualization and storage, the raw signal was digitally separated into local field potentials (LFP, 0.075–300 Hz; down-sampled to 1,375 Hz) and spike activity (SPK, 300– 9,000 Hz; sampling rate 44 kHz). Single-unit activity was detected and sorted online using amplitude thresholding and template-based waveform matching, with a predefined maximal deviation criterion between threshold-crossing waveforms and the template shapes. Up to four spike templates were defined per electrode. All data streams were time-synchronized and acquired using the AlphaLab SnR system (Alpha Omega Engineering, Nof Hagalil, Israel).

### Experimental procedures – MPTP administration and rehabilitation

The NHPs were treated with MPTP (1-methyl-4-phenyl-1,2,3,6-tetrahydropyridine)(*68*) after finishing recording under healthy conditions. Animals received five intramuscular (IM) injections of MPTP hydrochloride (0.35 mg/kg per injection; Sigma, Israel) over four consecutive days, including two injections on the first day. All procedures were conducted under ketamine sedation (10 mg/kg per injection, IM) in a negative-pressure isolation room. The NHPs were returned to their shared animal facility room 72 h after the last MPTP injection. They were kept in a separate space covered with a soft mattress to prevent pressure sores. Their peers still could interact and communicate with them through the enclosure. The NHP Parkinson’s score(*69*, *70*) was evaluated (1-2 weeks) before and (during all recording days) after MPTP. After MPTP, the NHPs were cared for intensively(*71*). They were fed twice daily, seven days per week, with a high-calorie nutritional shake (Ensure-Plus, Abbott) administered via a pediatric nasogastric tube. Dopamine replacement therapy (Dopicar; 250 mg levodopa/25 mg carbidopa per tablet) was given with the morning feeding on weekdays (10-12 h before afternoon/night recordings) and twice daily on weekends. To improve the therapeutic effect, dopamine replacement therapy was administered 15-30 minutes before feeding.

The recording protocol in the Parkinsonian state was the same as that in the healthy state, except for task performance. They were treated and cared for with an intensive rehabilitation protocol after the completion of the experimental protocol(*71*). All surgical attachments were removed from our experimental animals when their health condition allowed the surgical and anesthesia procedures. Monkey Md recovered and moved to the Israel Primate Sanctuary (www.ipsf.org.il). Monkey Wh did not recover and was euthanized to minimize suffering after discussions with the veterinary team and the ethical committee.

### Signal analysis – Polysomnography

The initial sleep staging was performed using a semi-automatic algorithm that clustered non-overlapping 10-s segments. Vigilance states, i.e., wakefulness, NREM sleep, REM sleep, and ambiguous/unclassified states, were identified based on the eye-open fraction, the root mean square (RMS) of the EMG signal, and the high-to-low EEG power ratio, defined as the average in the 15-25 Hz band divided by the average power in the 0.1-7 Hz band. The initial sleep classification was further refined based on the eye-open percentage, EMG activity, and the saccade frequency of the right eye. The processing of EMG and eye-coil signals, as well as the procedures for sleep classification, were previously described(*48*). Polysomnography examples before MPTP and during the MPTP-I and MPTP-II phases are shown in Figure 1b. The fractions of wakefulness, NREM, and REM sleep were calculated for each recording day. The fractions of those three states were normalized to their corresponding results before MPTP. The probabilities of vigilance states were also calculated.

To observe the effect of MPTP on REM sleep, we also analysed REM frequency (the number of REM per hour or per night), REM duration (the time of REM per hour or per night), latency to the 1^st^ REM episode (defined as latency from screen blackout to the onset of the first REM episode), first REM duration, last REM duration, the ratio of the last REM duration and the first REM duration (last REM/first REM). Finally, EMG RMS across vigilance states before and after MPTP was calculated and normalized by the average EMG RMS of the corresponding sleep states before MPTP.

### Signal analysis – fast eye movements (saccades)

Under healthy conditions, eye-coil signals were converted from voltage to angular position (degrees) using calibration data obtained on the same day (for more details, see Liu et al., 2026(*48*)). The calibration parameters obtained in the week prior to the MPTP administration were used for the MPTP recording.

Eye movements were classified as fast (saccades) and slow(drifts) (Fig. 2a-left). Examples of fast eye movements during wakefulness and REM sleep are shown in Figure 2a-right. In this study, we focused only on the fast eye movements. Saccades with amplitudes between 1 and 20 degrees and durations shorter than 200 milliseconds were included. To assess how MPTP affects eye movements across vigilance states, we calculated saccade frequency, amplitude, duration, and velocity.

Each saccade was represented as a binary (0/1) event occurring at the beginning of the saccade (saccade trains). We calculated the autocorrelation function of saccade trains and the coefficient of variance of the inter-saccade intervals (CV-ISacI). The autocorrelation histogram of saccade trains was calculated using the built-in function of MATLAB 2025b (xcorr, the correlation coefficient method). The mean value of each 10-s saccade train was subtracted to show the negative and positive correlation values. The time resolution of the saccade autocorrelation histogram was set to 0.1 seconds, and the lag range was from −2 to 2 s. The inter-saccade interval (ISacI) was defined as the duration between the first and second fast eye movements. CV-ISacI was defined as the standard deviation of ISacI divided by its mean (STD(ISacI)/mean(ISacI)). CV-ISacI was calculated for each 10-second epoch and averaged over the whole recording session.

### Signal analysis – filter and down-sampling of EMG, EEG, LFP, and SPK signals

The 50 Hz power artifact and its harmonics (100 Hz, 150 Hz, and 200 Hz) of EMG, EEG, and LFP were removed by a second-order IIR notch filter. Spiking activity (300-9000 Hz) was processed using an IIR comb-notch filter comprising 881 notches (44,000/50 + 1), designed to remove 50 Hz line noise and its harmonics, given a sampling rate of 44,000 Hz. The filter quality factor (Q) was set to 35, yielding a highly selective notch response. The filtered signals were then rectified by taking the absolute value, and the mean of the rectified signals was subtracted to obtain multi-unit activity (MUA).

Subsequently, the EEG, LFP, and MUA were filtered using a low-pass Butterworth filter with a 210 Hz passband frequency, 260 Hz stopband frequency, 1dB passband ripple, and 5 dB stopband attenuation. In the present study, EMG signals were digitally bandpass-filtered offline in the 10-500 Hz range, with stopbands at 0-5 Hz and 520-1375 Hz. The filtered EMG was used to calculate its RMS (root-mean-square) for each 10-s epoch. To minimize phase distortion, forward–backward filtering was applied, resulting in zero-phase filtering; this approach was used for all signals processed offline. Finally, the filtered EEG and MUA were down-sampled to 1/2 and 1/32 of their original sampling rate (resulting in 1375 Hz) by averaging every 2 and 32 consecutive samples, respectively. The DC (direct current, frequency = 0Hz) level was minimized by subtracting the average value of each signal epoch.

### Signal analysis – FOOOF analysis

The spectrograms of EEG, LFP, and spiking activities were separated into aperiodic and periodic components using fitting oscillations & one over f (FOOOF) algorithm, following in our previous study(*16*). EEG and LFP PSD were fitted with the following settings: peak_width_limits = [0.8, 12], peak_width_limits_per = [0.02, 0], max_n_peaks = 6, min_peak_height = 0.05, peak_threshold = 2, aperiodic_mode = “fixed”.

The envelope of the spiking activity (SPK) was calculated by the absolute operator. The parameters for SPK fitting were peak_width_limits = [0.5, 12], peak_width_limits_per = [0.02, 0], max_n_peaks = 8, min_peak_height = 0.05, peak_threshold = 2, aperiodic_mode = “fixed”. The fitting frequency range was from 3 to 40 Hz to avoid contamination by low-frequency artifacts from heart rate, breathing, pulsation of cerebrospinal fluid, or other sources, and to minimize inaccurate fitting across a broad frequency range.

The raw and periodic spectrograms in each state (wakefulness, NREM sleep, and REM sleep) were temporally normalized into 40 bins by averaging the power values within each time bin. Both aperiodic and periodic components were demonstrated in the reconstructed PSD. We also showed both components in aperiodic parameters (i.e., offset and exponent) and periodic parameters (i.e., center frequency and peak power). Finally, we also calculated the area under the curve of the periodic activity.

### Signal analysis – aligning power spectral density, beta peak power, saccade frequency, and EMG RMS to NREM-to-REM sleep transition

From the FOOOF analysis, we obtained the raw and periodic PSD and the beta peak power for each 10-s epoch, which are consistent with the sleep-state segments. The saccade frequency (number of saccades per second) and EMG RMS were calculated for each sleep 10-s epoch. Those variables were aligned to the first epoch of REM sleep. There were 30 NREM and 12 REM epochs before and after this sleep transition.

### Signal analysis – time-frequency domain spectrogram density of signals aligned to saccades

Time-domain EEG, LFP, and SPK activities were aligned with fast eye movements (saccades). The length of time-domain signals was 5 seconds before and after saccades, and these aligned signals were used to compute time-frequency spectrograms using a continuous wavelet transform with the analytic Morse wavelet and 1,375 Hz sampling rate (cwt, MATLAB R2025b, MathWorks). The continuous wavelet transform returned a matrix of complex values. We used the absolute value of these complex numbers. The time-frequency spectrograms beyond ±2 s and below 0.9 Hz are likely contaminated by edge effects. To ensure reliable power, only aligned spectrograms within the 0.9-200 Hz frequency range and 2 seconds before and after the saccade are shown.

The wavelet transforms provided frequency-dependent temporal resolution, with finer frequency resolution at lower frequencies and coarser frequency resolution at higher frequencies. Because the frequency bins of the continuous wavelet transform were not linearly spaced, the wavelet power values were normalized by the local frequency interval, computed as the gradient of the frequency vector. This procedure compensated for variable frequency resolution and approximated a linearly spaced spectral representation. Finally, they were normalized by frequency (calculating the power percentage of total power across the frequency range for each frequency at a given time point) and then z-score normalized relative to their baseline (from 2-1.5 s before the eye movement). To reduce the 3-D (time, frequency, and power) to 2-D (time and power), we averaged them within specific frequency ranges, e.g., the beta and theta-alpha bands.

### Statistical analysis and reproducibility

Statistical analyses were performed in MATLAB (R2025b, MathWorks). Population data are presented as the mean ± SEM (standard error of the mean). For comparison between independent samples, two-tailed Wilcoxon rank-sum tests were used; paired comparisons were assessed using two-tailed Wilcoxon signed-rank tests. Statistical significance was defined as *P* < 0.05 (two-tailed). When applicable, *P* values were adjusted for multiple comparisons using the Bonferroni correction. The *n* definition of sample size is clarified in the corresponding figure legend.

## Acknowledgements

The authors would like to thank Uri Werner-Reiss, PhD, for his valuable support with the surgical procedures and all aspects of the NHP care; Tamar Ravins Yaish, DMD, and the HUJI-ELSC animal facility team for their assistance with the NHP care. We acknowledge the use of large language model (LLM, ChatGPT-Go) tools for linguistic editing to improve the clarity and grammar of this manuscript; no scientific content was generated by these tools. This study is supported by grants from the ISF Breakthrough Research program (Grant No.: 1738/22) and the Collaborative Research Center TRR295, Germany (Project number 424778381) to HB.

## Author contributions

J.G. and H.B. conceived the research and designed the experiments. Z.I., D.W., and A.R. performed the surgical procedures. X.L. and J.G. supported the surgical procedure. They also performed experiments, including electrophysiological and behavioral recordings, analyzed the data, and conducted the statistical analysis. X.L., J.G., and H.B. prepared the figures and wrote the manuscript. H.B. supervised the work. All authors read and approved the final manuscript.

## Data availability

All data supporting the findings of this study are available within the manuscript as Supplementary Data. Other data will be available from the corresponding authors upon reasonable request.

## Code availability

We provided the code for the figures of this manuscript as a Supplementary File. MATLAB code will also be available from the corresponding authors upon reasonable request. Please note that the code used in this study was developed by the researchers for data analysis and visualization. It is intended for research purposes and may not meet professional coding standards.

## Correspondence and requests for materials should be addressed to

Xiaowei Liu, Jing Guang, or Hagai Bergman.

## References

1. C. Hammond, H. Bergman, P. Brown, Pathological synchronization in Parkinson’s disease: networks, models and treatments. Trends Neurosci. 30, 357–364 (2007).

2. E. Aserinsky, N. Kleitman, Regularly Occurring Periods of Eye Motility, and Concomitant Phenomena, During Sleep. Science 118, 273–274 (1953).

3. M. Jouvet, Neurophysiology of the states of sleep. Physiol. Rev. 47, 117–177 (1967).

4. P.-H. Luppi, J. Malcey, A. Chancel, B. Duval, S. Cabrera, P. Fort, Neuronal network controlling REM sleep. J. Sleep Res. 34, e14266 (2025).

5. C. H. Schenck, S. R. Bundlie, M. G. Ettinger, M. W. Mahowald, Chronic Behavioral Disorders of Human REM Sleep: A New Category of Parasomnia. Sleep 9, 293–308 (1986).

6. I. Arnulf, REM sleep behavior disorder: Motor manifestations and pathophysiology. Mov. Disord. 27, 677–689 (2012).

7. R. B. Postuma, A. Iranzo, M. Hu, B. Högl, B. F. Boeve, R. Manni, W. H. Oertel, I. Arnulf, L. Ferini-Strambi, M. Puligheddu, E. Antelmi, V. Cochen De Cock, D. Arnaldi, B. Mollenhauer, A. Videnovic, K. Sonka, K.-Y. Jung, D. Kunz, Y. Dauvilliers, F. Provini, S. J. Lewis, J. Buskova, M. Pavlova, A. Heidbreder, J. Y. Montplaisir, J. Santamaria, T. R. Barber, A. Stefani, E. K. St. Louis, M. Terzaghi, A. Janzen, S. Leu-Semenescu, G. Plazzi, F. Nobili, F. Sixel-Doering, P. Dusek, F. Bes, P. Cortelli, K. Ehgoetz Martens, J.-F. Gagnon, C. Gaig, M. Zucconi, C. Trenkwalder, Z. Gan-Or, C. Lo, M. Rolinski, P. Mahlknecht, E. Holzknecht, A. R. Boeve, L. N. Teigen, G. Toscano, G. Mayer, S. Morbelli, B. Dawson, A. Pelletier, Risk and predictors of dementia and parkinsonism in idiopathic REM sleep behaviour disorder: a multicentre study. Brain 142, 744–759 (2019).

8. V. C. De Cock, M. Vidailhet, S. Leu, A. Texeira, E. Apartis, A. Elbaz, E. Roze, J. C. Willer, J. P. Derenne, Y. Agid, I. Arnulf, Restoration of normal motor control in Parkinson’s disease during REM sleep. Brain 130, 450–456 (2007).

9. A. D. Mizrahi-Kliger, A. Kaplan, Z. Israel, M. Deffains, H. Bergman, Basal ganglia beta oscillations during sleep underlie Parkinsonian insomnia. Proc. Natl. Acad. Sci. 117, 17359–17368 (2020).

10. Z. Yin, R. Ma, Q. An, Y. Xu, Y. Gan, G. Zhu, Y. Jiang, N. Zhang, A. Yang, F. Meng, A. A. Kühn, H. Bergman, W.-J. Neumann, J. Zhang, Pathological pallidal beta activity in Parkinson’s disease is sustained during sleep and associated with sleep disturbance. Nat. Commun. 14, 5434 (2023).

11. W. Dement, The Effect of Dream Deprivation: The need for a certain amount of dreaming each night is suggested by recent experiments. Science 131, 1705–1707 (1960).

12. G. W. Vogel, A Review of REM Sleep Deprivation. Arch. Gen. Psychiatry 32, 749 (1975).

13. D. A. Robinson, “Is the oculomotor system a cartoon of motor control?” in Progress in Brain Research (Elsevier, 1986; https://linkinghub.elsevier.com/retrieve/pii/S0079612308634354) vol. 64, pp. 411–417.

14. D. A. Robinson, Oculomotor unit behavior in the monkey. J. Neurophysiol. 33, 393–403 (1970).

15. T. Donoghue, M. Haller, E. J. Peterson, P. Varma, P. Sebastian, R. Gao, T. Noto, A. H. Lara, J. D. Wallis, R. T. Knight, A. Shestyuk, B. Voytek, Parameterizing neural power spectra into periodic and aperiodic components. Nat. Neurosci. 23, 1655–1665 (2020).

16. X. Liu, J. Guang, S. Glowinsky, H. Abadi, D. Arkadir, E. Linetsky, M. Abu Snineh, J. F. León, Z. Israel, W. Wang, H. Bergman, Subthalamic nucleus input-output dynamics are correlated with Parkinson’s burden and treatment efficacy. Npj Park. Dis. 10, 117 (2024).

17. E. Urrestarazu, J. Iriarte, M. Alegre, P. Clavero, M. C. Rodríguez-Oroz, J. Guridi, J. A. Obeso, J. Artieda, Beta activity in the subthalamic nucleus during sleep in patients with Parkinson’s disease. Mov. Disord. 24, 254–260 (2009).

18. A. Yugeta, W. D. Hutchison, C. Hamani, U. Saha, A. M. Lozano, M. Hodaie, E. Moro, R. Chen, Modulation of Beta Oscillations in the Subthalamic Nucleus with Prosaccades and Antisaccades in Parkinson’s Disease. J. Neurosci. 33, 6895–6904 (2013).

19. G. Pfurtscheller, C. Neuper, Motor imagery activates primary sensorimotor area in humans. Neurosci. Lett. 239, 65–68 (1997).

20. A. K. Engel, P. Fries, Beta-band oscillations — signalling the status quo? Curr. Opin. Neurobiol. 20, 156–165 (2010).

21. A. R. Aron, R. A. Poldrack, Cortical and Subcortical Contributions to Stop Signal Response Inhibition: Role of the Subthalamic Nucleus. J. Neurosci. 26, 2424–2433 (2006).

22. Y. Bao, C. Gan, Z. Chen, Z. Qi, Z. Meng, F. Yue, Quantification of Non-Motor Symptoms in Parkinsonian Cynomolgus Monkeys. Brain Sci. 13, 1153 (2023).

23. A. Davin, S. Chabardès, H. Belaid, D. Fagret, L. Djaileb, Y. Dauvilliers, O. David, N. Torres-Martinez, B. Piallat, Early onset of sleep/wake disturbances in a progressive macaque model of Parkinson’s disease. Sci. Rep. 12, 17499 (2022).

24. H. Belaid, J. Adrien, C. Karachi, E. C. Hirsch, C. François, Effect of melatonin on sleep disorders in a monkey model of Parkinson’s disease. Sleep Med. 16, 1245–1251 (2015).

25. K. Fifel, J. Vezoli, K. Dzahini, B. Claustrat, V. Leviel, H. Kennedy, E. Procyk, O. Dkhissi-Benyahya, C. Gronfier, H. M. Cooper, Alteration of Daily and Circadian Rhythms following Dopamine Depletion in MPTP Treated Non-Human Primates. PLoS ONE 9, e86240 (2014).

26. P. S. Verhave, M. J. Jongsma, R. M. Van Den Berg, J. C. Vis, R. A. P. Vanwersch, A. B. Smit, E. J. W. Van Someren, I. H. C. H. M. Philippens, REM Sleep Behavior Disorder in the Marmoset MPTP Model of Early Parkinson Disease. Sleep 34, 1119–1125 (2011).

27. H. Belaid, J. Adrien, E. Laffrat, D. Tande, C. Karachi, D. Grabli, I. Arnulf, S. D. Clark, X. Drouot, E. C. Hirsch, C. Francois, Sleep Disorders in Parkinsonian Macaques: Effects of L-Dopa Treatment and Pedunculopontine Nucleus Lesion. J. Neurosci. 34, 9124–9133 (2014).

28. Q. Barraud, V. Lambrecq, C. Forni, S. McGuire, M. Hill, B. Bioulac, E. Balzamo, E. Bezard, F. Tison, I. Ghorayeb, Sleep disorders in Parkinson’s disease: The contribution of the MPTP non-human primate model. Exp. Neurol. 219, 574–582 (2009).

29. H. Almirall, I. Pigarev, M. D. De La Calzada, M. Pigareva, M. T. Herrero, T. Sagales, Nocturnal sleep structure and temperature slope in MPTP treated monkeys. J. Neural Transm. 106, 1125–1134 (1999).

30. K. Pungor, A. Hajnal, K. A. Kékesi, G. Juhász, Paradoxical sleep deprivatory effect of a single low dose of MPTP which did not produce dopaminergic cell loss. Exp. Brain Res. 95 (1993).

31. A. McCarthy, K. Wafford, E. Shanks, M. Ligocki, D. M. Edgar, D.-J. Dijk, REM sleep homeostasis in the absence of REM sleep: Effects of antidepressants. Neuropharmacology 108, 415–425 (2016).

32. A. L. Sharpley, P. J. Cowen, Effect of pharmacologic treatments on the sleep of depressed patients. Biol. Psychiatry 37, 85–98 (1995).

33. H.-P. Landolt, E. B. Raimo, B. J. Schnierow, J. R. Kelsoe, M. H. Rapaport, J. C. Gillin, Sleep and Sleep Electroencephalogram in Depressed Patients Treated With Phenelzine. Arch. Gen. Psychiatry 58, 268 (2001).

34. R. J. Wyatt, Total Prolonged Drug-Induced REM Sleep Suppression in Anxious-Depressed Patients. Arch. Gen. Psychiatry 24, 145 (1971).

35. A. Dagay, S. Katzav, N. Elisha, J. Volkov, R. Tauman, N. Giladi, J. M. Hausdorff, A. Mirelman, J. Zitser, REM density in Parkinson’s disease: association with motor, cognitive, autonomic function, and dopaminergic medication. Npj Park. Dis. 11, 211 (2025).

36. Y. Zhang, R. Ren, L. D. Sanford, L. Yang, J. Zhou, L. Tan, T. Li, J. Zhang, Y.-K. Wing, J. Shi, L. Lu, X. Tang, Sleep in Parkinson’s disease: A systematic review and meta-analysis of polysomnographic findings. Sleep Med. Rev. 51, 101281 (2020).

37. E. Aserinsky, A. L. Joan, M. E. Mack, S. P. Tzankoff, E. Hurn, Comparison of Eye Motion in Wakefulness and REM Sleep. Psychophysiology 22, 1–10 (1985).

38. J. R. Hotson, E. B. Langston, J. W. Langston, Saccade responses to dopamine in human MPTP-induced parkinsonism. Ann. Neurol. 20, 456–463 (1986).

39. H. Slovin, M. Abeles, E. Vaadia, I. Haalman, Y. Prut, H. Bergman, Frontal Cognitive Impairments and Saccadic Deficits in Low-Dose MPTP-Treated Monkeys. J. Neurophysiol. 81, 858–874 (1999).

40. Z. Yin, T. Yuan, A. Yang, Y. Xu, G. Zhu, Q. An, R. Ma, Y. Gan, L. Shi, Y. Bai, N. Zhang, C. Wang, Y. Jiang, F. Meng, W.-J. Neumann, H. Tan, J.-G. Zhang, Contribution of basal ganglia activity to REM sleep disorder in Parkinson’s disease. J. Neurol. Neurosurg. Psychiatry, jnnp-2023-332014 (2024).

41. A. K. Verma, S. F. Acosta Lenis, J. E. Aman, D. E. Sanabria, J. Wang, A. Pearson, M. Hill, R. Patriat, L. E. Schrock, S. E. Cooper, M. C. Park, N. Harel, M. J. Howell, C. D. MacKinnon, J. L. Vitek, L. A. Johnson, Basal ganglia engagement during REM sleep movements in Parkinson’s disease. Npj Park. Dis. 8, 116 (2022).

42. M. Hackius, E. Werth, O. Sürücü, C. R. Baumann, L. L. Imbach, Electrophysiological Evidence for Alternative Motor Networks in REM Sleep Behavior Disorder. J. Neurosci. 36, 11795–11800 (2016).

43. R. Ferri, F. Rundo, A. Silvani, M. Zucconi, O. Bruni, L. Ferini-Strambi, G. Plazzi, M. Manconi, REM Sleep EEG Instability in REM Sleep Behavior Disorder and Clonazepam Effects. Sleep 40 (2017).

44. M. M. Hoehn, M. D. Yahr, Parkinsonism: onset, progression and mortality. Neurology 17, 427– 442 (1967).

45. Y. E. Kim, H. J. Yang, J. Y. Yun, H.-J. Kim, J.-Y. Lee, B. S. Jeon, REM sleep behavior disorder in Parkinson disease: Association with abnormal ocular motor findings. Parkinsonism Relat. Disord. 20, 444–446 (2014).

46. A. Cicolin, L. Lopiano, M. Zibetti, E. Torre, A. Tavella, G. Guastamacchia, A. Terreni, G. Makrydakis, E. Fattori, M. M. Lanotte, B. Bergamasco, R. Mutani, Effects of deep brain stimulation of the subthalamic nucleus on sleep architecture in parkinsonian patients. Sleep Med. 5, 207–210 (2004).

47. N. Hjort, K. Østergaard, E. Dupont, Improvement of sleep quality in patients with advanced Parkinson’s disease treated with deep brain stimulation of the subthalamic nucleus. Mov. Disord. 19, 196–199 (2004).

48. X. Liu, J. Guang, Z. Israel, D. Wajnsztajn, A. Raz, H. Bergman, Entrained cortical delta–spindle activity, not periodic synchrony, prevents arousal by NREM thalamic bursts. *Commun*. Biol. 9, 285 (2026).

49. G. Pfurtscheller, A. Stancák, Ch. Neuper, Event-related synchronization (ERS) in the alpha band — an electrophysiological correlate of cortical idling: A review. Int. J. Psychophysiol. 24, 39–46 (1996).

50. L. Shirahige, M. Berenguer-Rocha, S. Mendonça, S. Rocha, M. C. Rodrigues, K. Monte-Silva, Quantitative Electroencephalography Characteristics for Parkinson’s Disease: A Systematic Review. J. Park. Dis. 10, 455–470 (2020).

51. C. Moisello, D. Blanco, J. Lin, P. Panday, S. P. Kelly, A. Quartarone, A. Di Rocco, C. Cirelli, G. Tononi, M. F. Ghilardi, Practice changes beta power at rest and its modulation during movement in healthy subjects but not in patients with Parkinson’s disease. Brain Behav. 5, e00374 (2015).

52. N. C. Rowland, C. De Hemptinne, N. C. Swann, S. Qasim, S. Miocinovic, J. L. Ostrem, R. T. Knight, P. A. Starr, Task-related activity in sensorimotor cortex in Parkinson’s disease and essential tremor: changes in beta and gamma bands. Front. Hum. Neurosci. 9 (2015).

53. M. R. C. Van Den Heuvel, E. E. H. Van Wegen, P. J. Beek, G. Kwakkel, A. Daffertshofer, Incongruent visual feedback during a postural task enhances cortical alpha and beta modulation in patients with Parkinson’s disease. Clin. Neurophysiol. 129, 1357–1365 (2018).

54. G. Tamás, I. Szirmai, L. Pálvölgyi, A. Takáts, A. Kamondi, Impairment of post-movement beta synchronisation in parkinson’s disease is related to laterality of tremor. Clin. Neurophysiol. 114, 614–623 (2003).

55. A. J. Baumgartner, C. A. Kushida, M. O. Summers, D. S. Kern, A. Abosch, J. A. Thompson, Basal Ganglia Local Field Potentials as a Potential Biomarker for Sleep Disturbance in Parkinson’s Disease. Front. Neurol. 12, 765203 (2021).

56. L. Guan, H. Yu, Y. Chen, C. Gong, H. Hao, Y. Guo, S. Xu, Y. Zhang, X. Yuan, G. Yin, J. Zhang, H. Tan, L. Li, Subthalamic γ Oscillation Underlying Rapid Eye Movement Sleep Abnormality in Parkinsonian Patients. Mov. Disord. 40, 456–467 (2025).

57. O. Manzanilla, M. Alegre, A. Horrillo-Maysonnial, E. Urrestarazu, M. Valencia, Cortical activation in REM behavior disorder mimics voluntary movement. An electroencephalography study. Clin. Neurophysiol. 166, 191–198 (2024).

58. J. H. Shin, J.-Y. Lee, Y.-K. Kim, E. J. Yoon, H. Kim, H. Nam, B. Jeon, Parkinson Disease-Related Brain Metabolic Patterns and Neurodegeneration in Isolated REM Sleep Behavior Disorder. Neurology 97, e378–e388 (2021).

59. V. D. Sharma, S. Sengupta, S. Chitnis, A. W. Amara, Deep Brain Stimulation and Sleep-Wake Disturbances in Parkinson Disease: A Review. Front. Neurol. 9, 697 (2018).

60. A. A. Kühn, D. Williams, A. Kupsch, P. Limousin, M. Hariz, G. Schneider, K. Yarrow, P. Brown, Event-related beta desynchronization in human subthalamic nucleus correlates with motor performance. Brain 127, 735–746 (2004).

61. S. Cascino, F. Luiso, L. Caffi, C. Palmisano, E. Contaldi, G. Pezzoli, I. U. Isaias, S. Bonvegna, Chronic adaptive deep brain stimulation in Parkinson’s disease: ADAPT-START findings and programming principles. Npj Park. Dis. 12, 85 (2026).

62. W. F. Asaad, S. A. Sheth, What’s the n? On sample size vs. subject number for brain-behavior neurophysiology and neuromodulation. Neuron 112, 2086–2090 (2024).

63. E. Psarou, C. Katsanevaki, E. Maris, P. Fries, Would You Agree If N Is Three? On Statistical Inference for Small N. J. Cogn. Neurosci. 38, 1078–1088 (2026).

64. Guide for the Care and Use of Laboratory Animals: Eighth Edition (National Academies Press, Washington, D.C., 2011; https://www.nationalacademies.org/publications/12910).

65. D. Valsky, K. T. Blackwell, I. Tamir, R. Eitan, H. Bergman, Z. Israel, Real-time machine learning classification of pallidal borders during deep brain stimulation surgery. J. Neural Eng. 17, 016021 (2020).

66. D. Valsky, O. Marmor-Levin, M. Deffains, R. Eitan, K. T. Blackwell, H. Bergman, Z. Israel, Stop! border ahead: Automatic detection of subthalamic exit during deep brain stimulation surgery. Mov. Disord. 32, 70–79 (2017).

67. S. Glowinsky, Z. Israel, S. Heymann, H. Bergman, Divide and Conquer: Automatic Detection of the Thalamus to Empower DBS Physiological Navigation to the Subthalamic Region. IEEE Trans. Neural Syst. Rehabil. Eng. 33, 2672–2683 (2025).

68. J. Guang, H. Baker, O. Ben-Yishay Nizri, S. Firman, U. Werner-Reiss, V. Kapuller, Z. Israel, H. Bergman, Toward asleep DBS: cortico-basal ganglia spectral and coherence activity during interleaved propofol/ketamine sedation mimics NREM/REM sleep activity. Npj Park. Dis. 7, 67 (2021).

69. C. Imbert, E. Bezard, S. Guitraud, T. Boraud, C. E. Gross, Comparison of eight clinical rating scales used for the assessment of MPTP-induced parkinsonism in the Macaque monkey. J. Neurosci. Methods 96, 71–76 (2000).

70. A. Benazzouz, T. Boraud, P. Dubédat, A. Boireau, J.-M. Stutzmann, C. Gross, Riluzole prevents MPTP-induced parkinsonism in the rhesus monkey: a pilot study. Eur. J. Pharmacol. 284, 299– 307 (1995).

71. T. Ravins Yaish, N. Eshkol Noy, R. Kalman, J. Guang, H. Baker Erdman, O. Ben-Yishay Nizri, S. Firman, X. Liu, M. Deffains, U. Werner-Reiss, G. Abourbeh, Z. Israel, H. Bergman, L. Iskhakova, Innovative care protocol successfully rehabilitates non-human primates after MPTP-induced parkinsonism: Preliminary evidence from a restricted cohort of African Green Monkeys (*Chlorocebus sabaeus*). Lab. Anim. 59, 523–529 (2025).

