## Supplementary material for "REM Sleep Disengagement of β Oscillations Permits Rapid Dream Movements in Parkinsonism": Figures S1-S7

**Title**

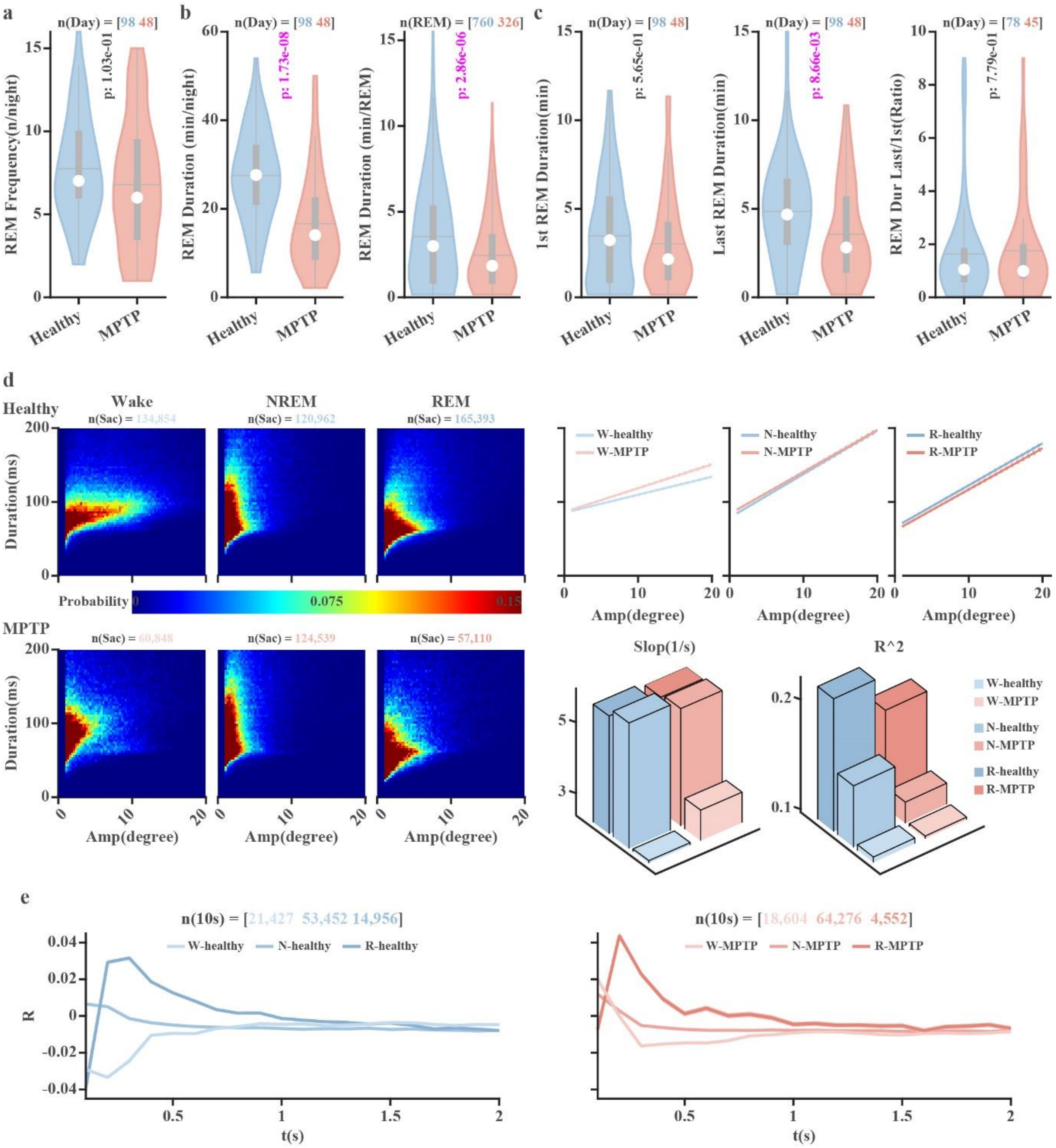


**Figure S1. MPTP affects REM sleep and saccade dynamics across vigilance states. a.** The number of REM sleep episodes per night before and after MPTP. **b.** Duration of REM sleep per night and the duration of a single REM sleep episode before and after MPTP. **c.** Duration of the 1st REM episode (left), the duration of the last REM episode (middle), and the ratio of the duration of the last REM episode to the duration of the first REM (right) before and after MPTP. Ratios of the last and 1st REM duration larger than 10 are removed. The violin plots in **a-c** show the mean (grey horizontal lines), median (white circles), 25th and 75th percentiles (the bottom and top edges of the grey boxes, respectively), and the spread of data (grey whiskers). **d.** The heat map of saccade duration and saccade amplitude (before and after MPTP, first and second row). The linear fittings of the amplitude-duration regression lines in corresponding vigilance states are shown in the right upper panel. The slopes and goodness-of-fit ($R^2$) of each linear fitting are shown in the right lower panel. **e.** Autocorrelation histograms of saccades across vigilance states before (left panel) and after (right panel) MPTP administration. The R values are shown as mean ± SEM (solid and shading). The number of units is shown in the title of each subplot. The statistical difference of variables between before and after MPTP in **a-c** subplots is calculated using the Wilcoxon rank sum test ($p<0.05$). The p-values indicating significant differences are marked in magenta; otherwise, they are black. Abbreviations and color coding as in Figure 1. Related to Figures 1 and 2.

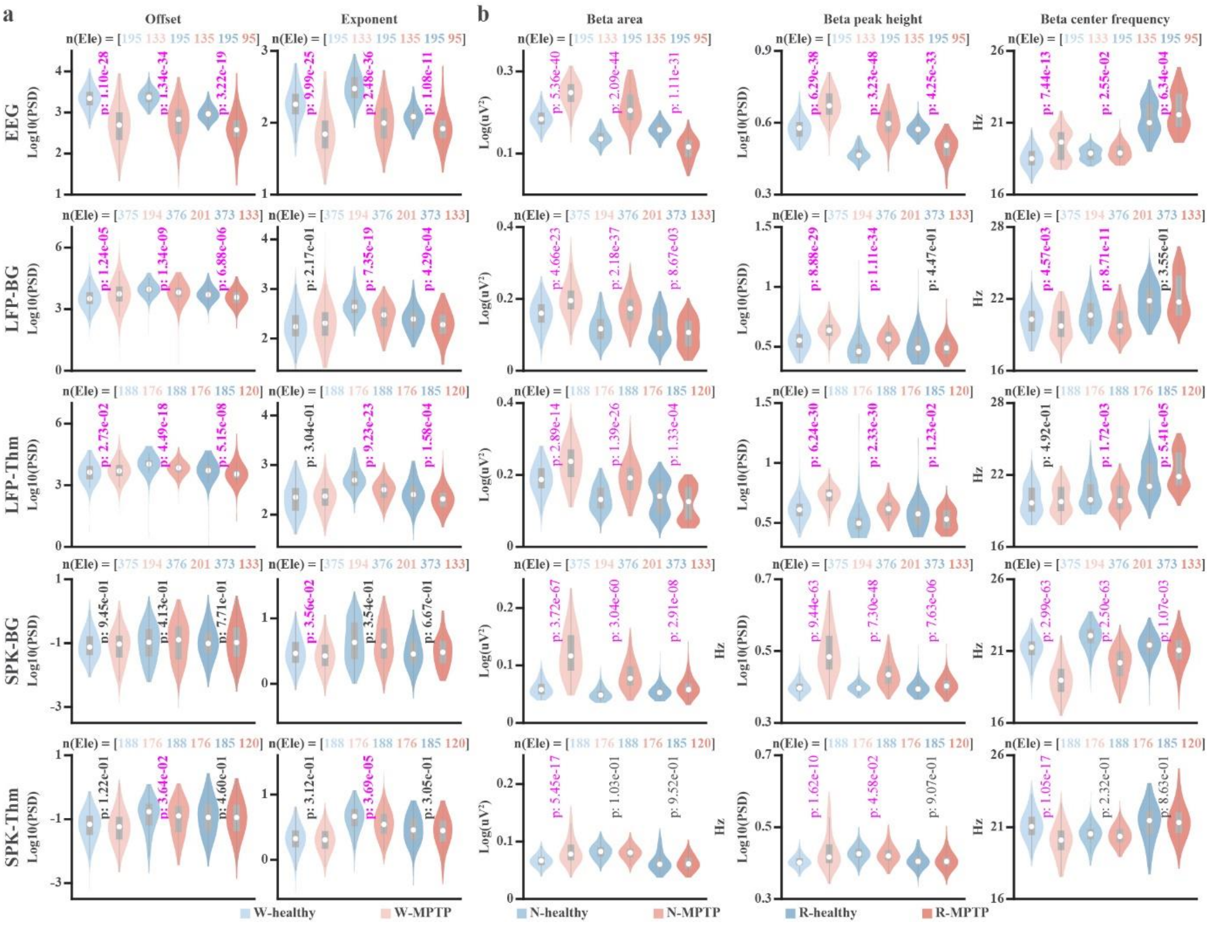


**Figure S2. The distribution and statistical information of parameters for the neuronal signals before and after MPTP administration across vigilance states. a.** The aperiodic parameters (offset and exponent) of frontal EEG, basal ganglia (BG), and thalamic (Thm) LFP and SPK activity before and after MPTP administration across vigilance states. **b.** The periodic beta area (column 1), peak power (column 2), and beta center frequency (column 3) before and after MPTP administration across vigilance states. The beta area is the summation of power in the beta frequency range (the summation of log10(beta power) * log10(frequency)). The violin plots show the median (white circles), 25th and 75th percentiles (the bottom and top edges of the grey boxes, respectively), and the spread of data (grey whiskers). The number of electrodes (Ele) is marked in the title of each subplot. The statistical difference in variables between before and after MPTP is calculated using the Wilcoxon rank-sum test ($p < 0.05$). The p-values indicating significant differences are marked in magenta; otherwise, they are black. Abbreviations and color coding are as in Figure 1. Related to Figures 4 and 5.

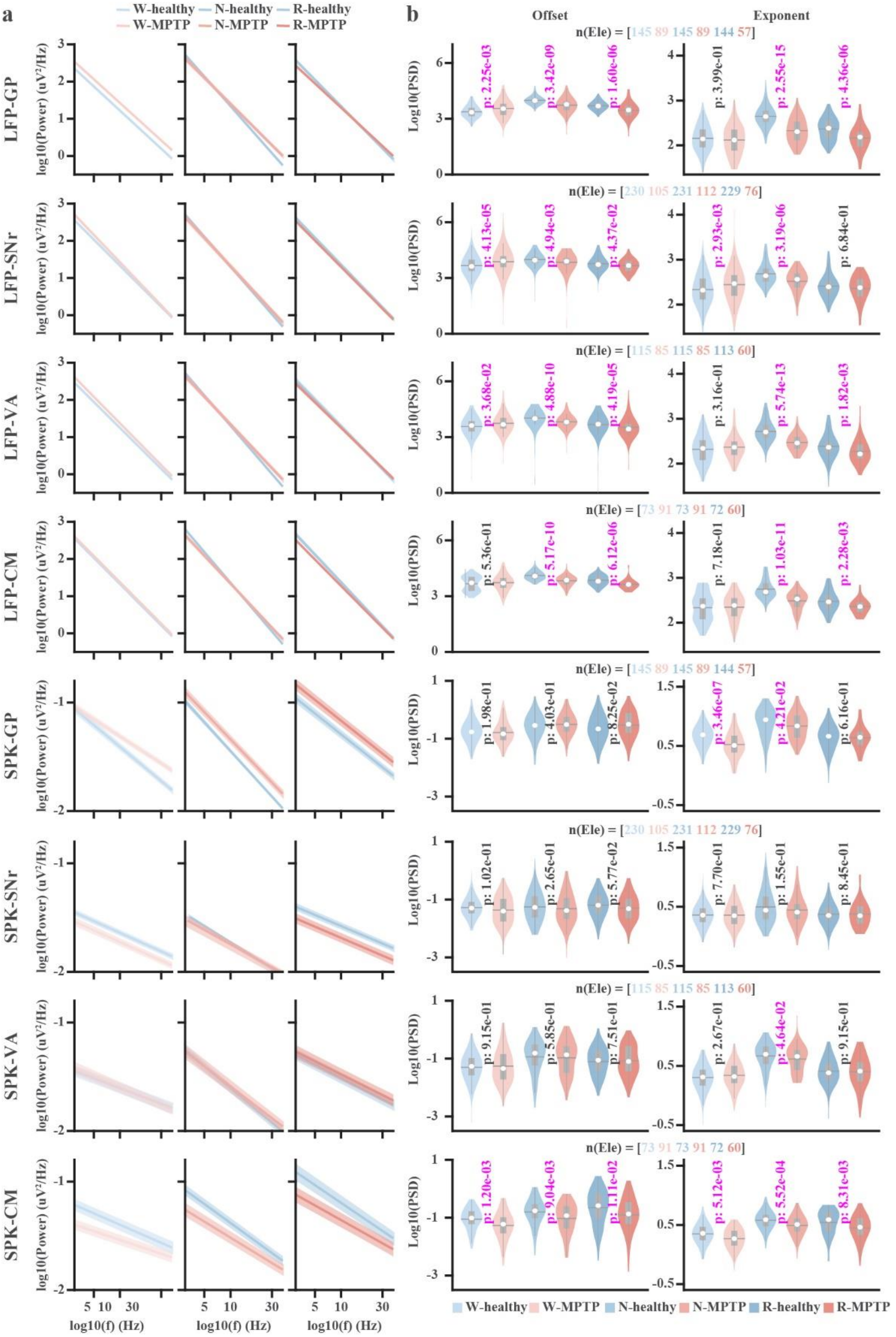

**Figure S3. Aperiodic LFP and spiking features before and after MPTP across vigilance states in distinct subcortical nuclei**. **a**. One-over-f fitting of aperiodic power of LFP and SPK before and after MPTP across vigilance states in the external segment of globus pallidus (GPe), substantia nigra pars reticulata (SNr), and the ventral anterior (VA) and centromedian (CM) of the thalamus. **b.** Violin plots of the distribution of the aperiodic parameters (offset and exponent) before and after MPTP across vigilance states in each corresponding subcortical nucleus. The data in **a** are shown as mean ± SEM. The number of electrodes (Ele) is the same as that revealed in the subtitle of **b.** The violin plots in **b** show the mean (grey horizontal lines), median (white circles), 25th and 75th percentiles (the bottom and top edges of the grey boxes, respectively), and the spread of data (grey whiskers). The statistical difference of aperiodic parameters between before and after MPTP in **b** is calculated using the Wilcoxon rank sum test ($p<0.05$). The p-values in **b** indicating significant differences are marked in magenta; otherwise, they are black. Abbreviations and color coding are as in Figure 1. Related to Figure 4.

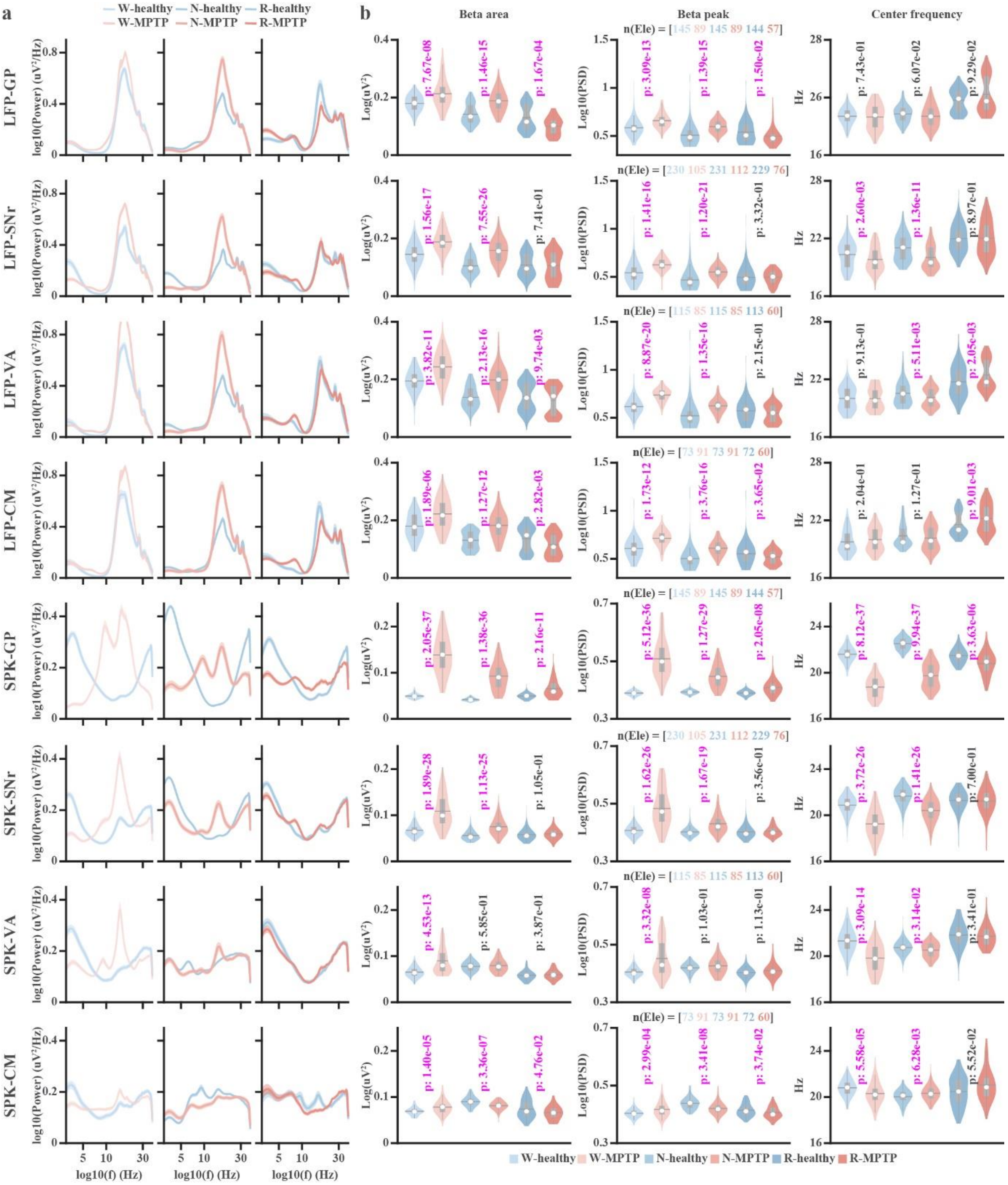


**Figure S4. Periodic LFP and spiking features before and after MPTP across vigilance states in distinct subcortical nuclei**. **a**. Gaussian fit of aperiodic power of LFP and SPK activity before and after MPTP across vigilance states in the external segment of globus pallidus (GPe), substantia nigra pars reticulata (SNr), and in the ventral anterior (VA) and centromedian (CM) of the thalamus. **b.** Violin plots of the distribution of the periodic beta area and periodic parameters (peak power and center frequency of beta oscillation) in each corresponding subcortical nucleus across vigilance states before and after MPTP. The data in **a** are shown as mean ± SEM. The number of electrodes (Ele) is the same as that revealed in the subtitle of **b.** The violin plots show the mean (grey horizontal lines), median (white circles), 25th and 75th percentiles (the bottom and top edges of the grey boxes, respectively), and the spread of data (grey whiskers). The statistical difference of periodic parameters between before and after MPTP in **b** is calculated using the Wilcoxon rank sum test ($p<0.05$). The p-values in **b** indicating significant differences are marked in magenta; otherwise, they are black. Abbreviations and color coding are as in Figure 1. Related to Figure 5.

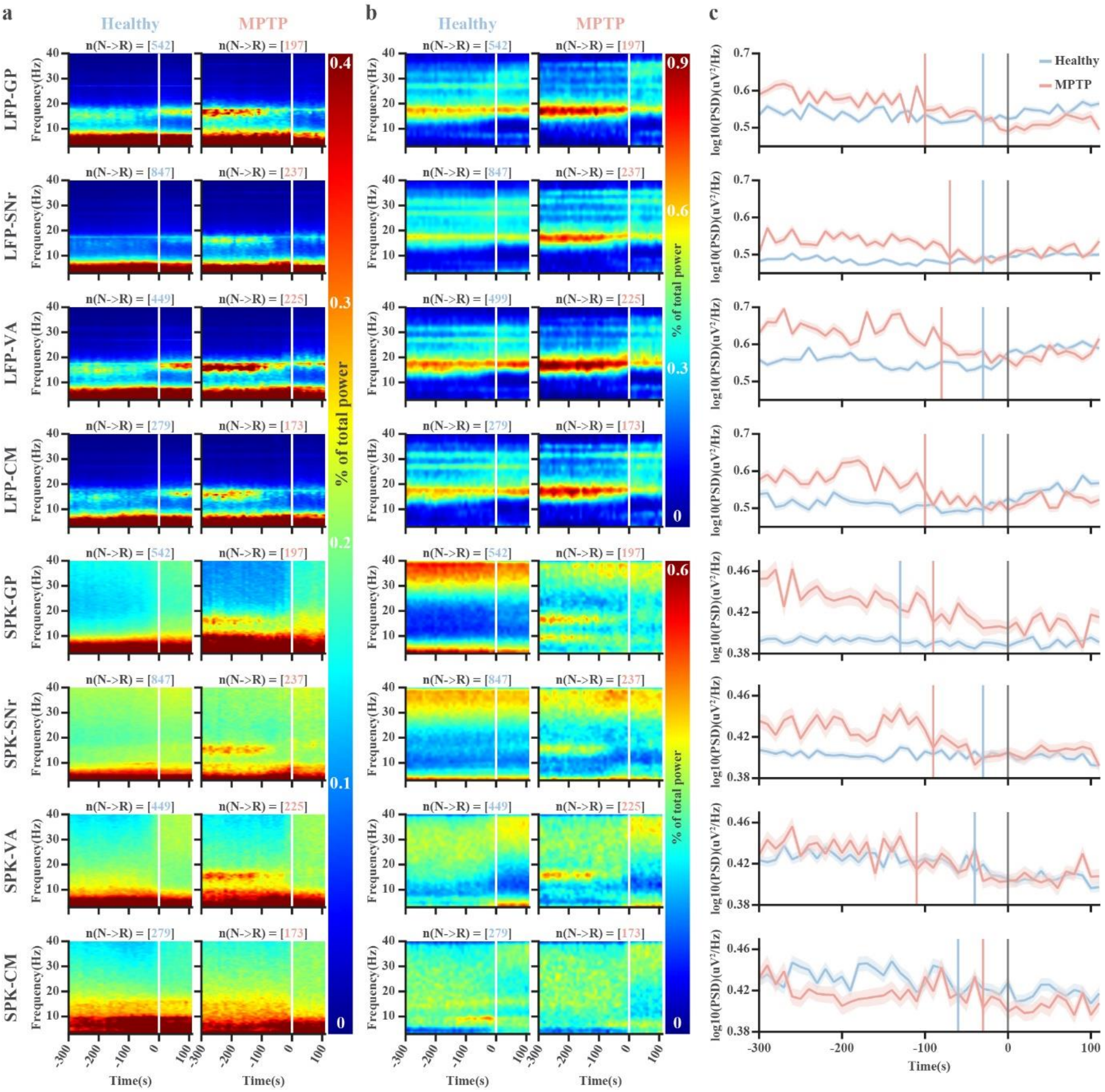


**Figure S5. Beta power changes across the transition from NREM to REM sleep in subcortical nuclei**. **a.** The raw spectrograms of LFP and SPK before and after MPTP in the external segment of globus pallidus (GPe), substantia nigra pars reticulata (SNr), and in the ventral anterior (VA) and centromedian (CM) nuclei of the thalamus are aligned to the NREM-to-REM sleep transition. **b.** The periodic spectrograms of LFP and SPK before and after MPTP in the GPe, SNr, VA, and CM nuclei are aligned to the NREM-to-REM sleep transition. The vertical white lines indicate the first epoch of REM sleep (time 0). There are 30 and 12 10-second NREM and REM segments before and after time 0, respectively. **c**. The corresponding peak power of beta oscillations aligned to the NREM-REM sleep transition (time=0, vertical grey lines). Blue/red vertical lines mark the change points of the beta power before and after MPTP. The number of NREM-to-REM sleep transitions in **a, b,** and **e** is shown in the title of the corresponding subplots. The number of NREM-to-REM sleep transitions in **c** is the same as that in **b.** The data in **c** are shown as mean ± SEM. Abbreviations and color coding are as in Figure 1. Related to Figure 6.

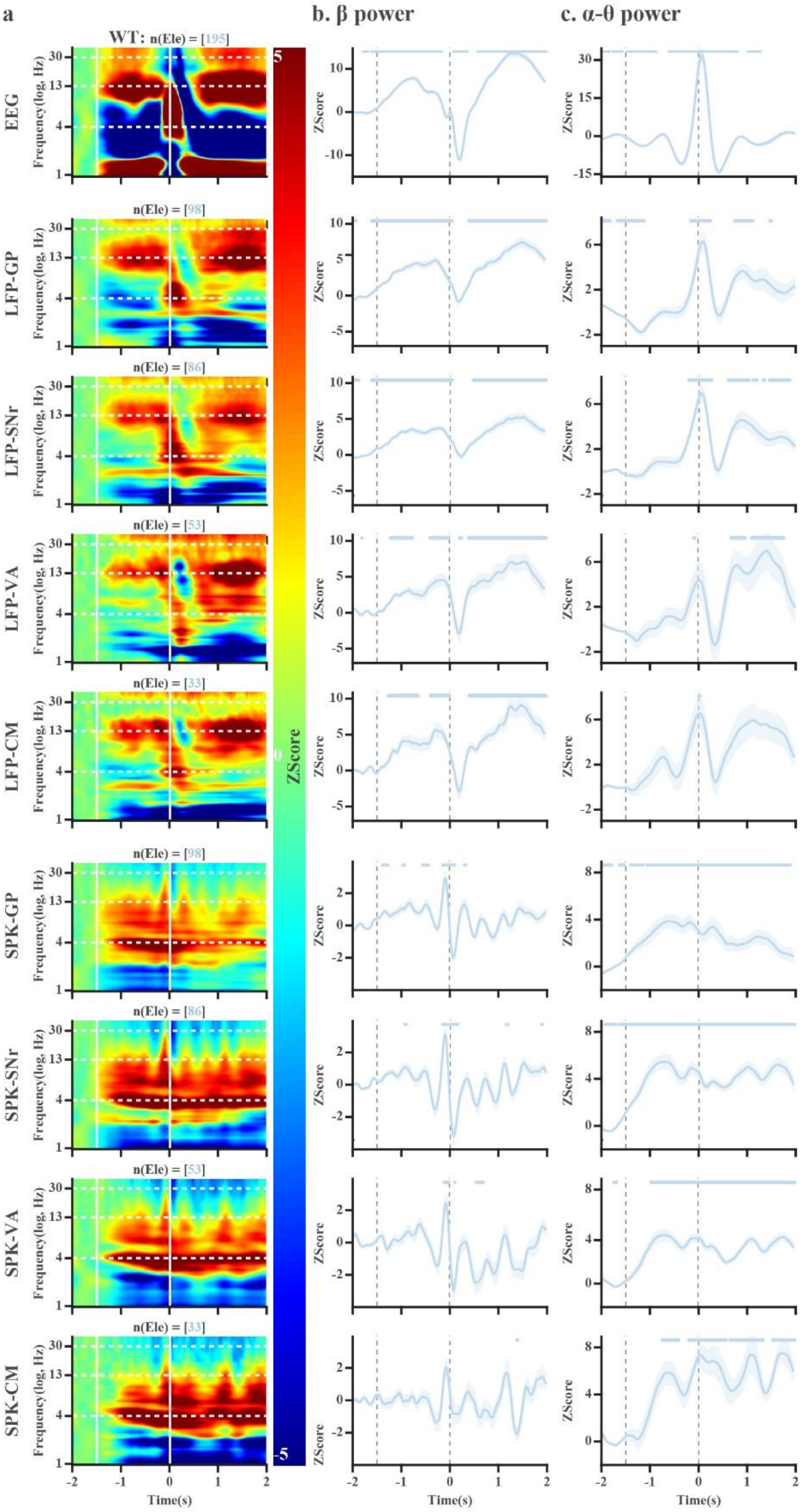


**Figure S6. Saccade-aligned EEG, LFP, and spiking spectrograms during task performance reveal beta desynchronization and theta-alpha synchronization before MPTP**. **a.** The spectrograms of frontal EEG, LFP and SPK in the external segment of globus pallidus (GPe), substantia nigra pars reticulata (SNr), and in the ventral anterior (VA) and centromedian (CM) thalamic nuclei are aligned to the voluntary (task) saccades. The first vertical white lines indicate the baseline (-2 to -1.5 seconds before time 0) period used for the z-score normalization. The second vertical line indicates saccade onset (time 0). The beta frequency range is shown between the two upper horizontal white dashed lines. The theta-alpha frequency range is shown between the 2nd and 3rd horizontal white dashed lines. The number of electrodes (Ele) is shown in the title of each subplot. **b.** The average power of neuronal activity in the beta frequency range. **c.** The average power of neuronal activity in the theta-alpha frequency range. In **b** and **c,** the number of electrodes is the same as that in **a**, and data are shown as mean ± SEM. The vertical grey dashed lines have the same meaning as the vertical white lines in **a**. The blue dots at the top of each axis indicate a significant difference from the baseline (from -2 to -1.5 seconds before the saccade). The Bonferroni-corrected Wilcoxon rank-sum test was used to assess differences in beta or theta-alpha power across lag time points ($p < 0.05/1375$) in **b** and **c**. Abbreviations and color coding are as in Figure 1. Related to Figure 7.

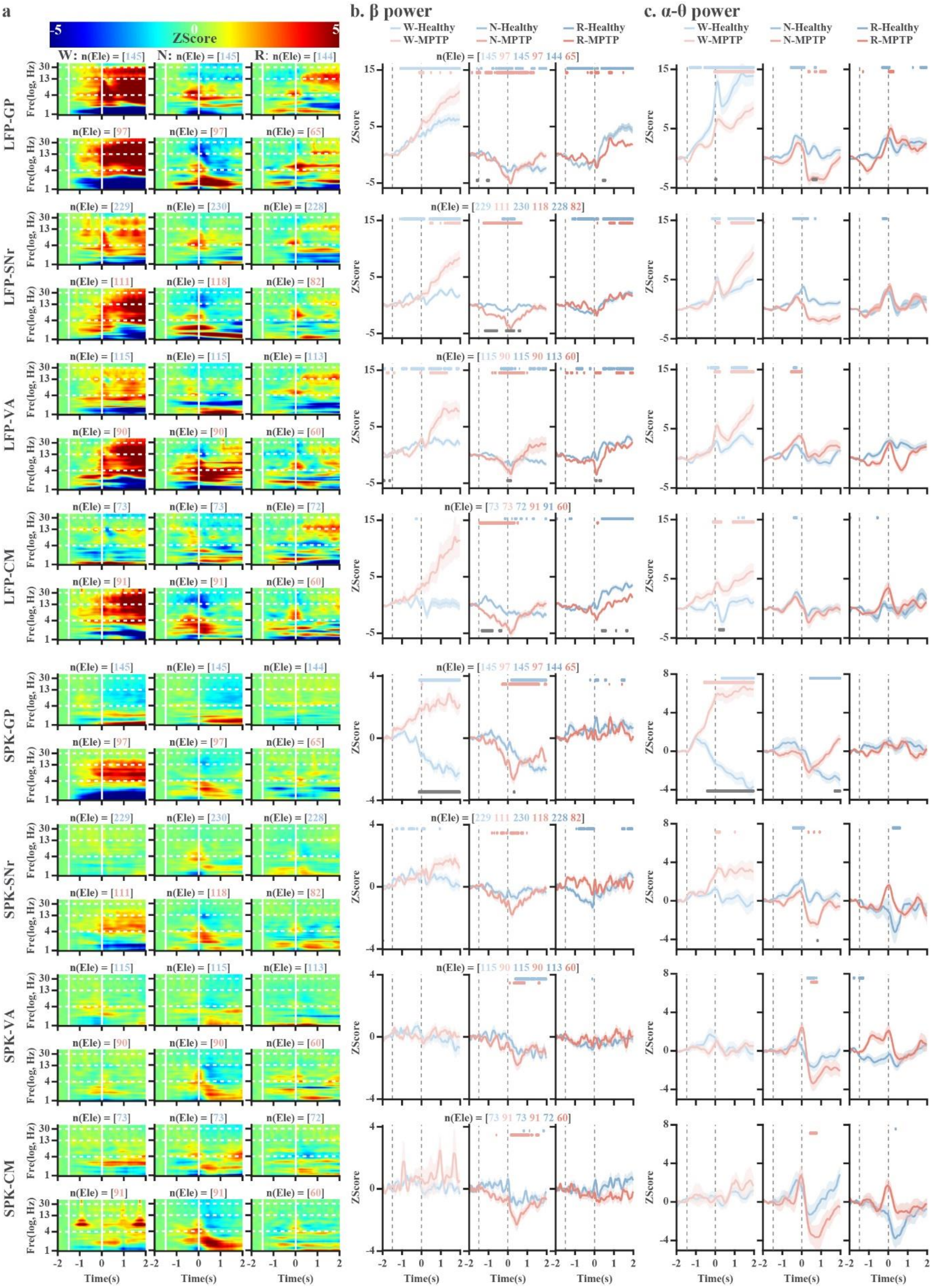


**Figure S7. Saccade-aligned EEG, LFP, and spiking spectrograms before and after MPTP across vigilance states in subcortical nuclei reveal state-dependent beta desynchronization and theta-alpha synchronization**. **a.** The spectrograms of LFP and SPK in the external segment of globus pallidus (GPe), substantia nigra pars reticulata (SNr), and in the ventral anterior (VA) and centromedian (CM) thalamic nuclei are aligned to fast eye movements (saccades). The number of electrodes (Ele) is shown in each subplot. The 1st vertical white line indicates the baseline (-2 to -1.5 seconds before time 0) used for the z-score normalization. The 2nd vertical line indicates saccades (time 0). The beta frequency range is shown between the upper horizontal white dashed lines. The theta-alpha frequency range is between the 2nd and 3rd horizontal white dashed lines. The Y-axis is frequency (Fre) on a log scale. **b.** The average power of the beta frequency range in **a**. **c.** The average power of the theta-alpha frequency range in **a**. In **b** and **c,** the number of electrodes is the same as that in **a**, and data are shown as mean ± SEM. The left and right vertical grey lines indicate the baseline of normalization and fast eye movement, respectively. The blue and pink dots at the top of each axis indicate significant differences from the baseline (between -2 and -1.5 seconds before saccade). The Bonferroni-corrected Wilcoxon rank-sum test was used to assess the difference in beta or theta-alpha power across different lag time points ($p < 0.05/1375$) in **b** and **c**. The grey dots at the bottom of the subplots indicate a significant difference in beta or theta-alpha power between before and after MPTP (Wilcoxon rank-sum test, $p<0.05$). Abbreviations and color coding are as in Figure 1. Related to Figure 7.
